# ITK deficiency protects against autoimmune pulmonary hemorrhage by promoting Amphiregulin-producing regulatory T cells

**DOI:** 10.64898/2026.05.01.722312

**Authors:** Mohammad Saddam Hossain, Ali Mobeen, Ibrahim H. Topkaya, Hui Xiong, Rosanne Trevail, Li Chen, Liye Suo, Mobin Karimi

**Affiliations:** Department of Microbiology and Immunology, SUNY Upstate Medical University, Syracuse, NY 13210; School of Medical Imaging, Nanchang Medical College, Nanchang, Jiangxi, Public Republic of China; Department of Pathology, SUNY Upstate Medical University, Syracuse, NY 13210

## Abstract

Pulmonary hemorrhage (PH) is a life-threatening manifestation of systemic autoimmunity caused by immune-mediated disruption of the alveolar capillary barrier. Despite high mortality, the molecular checkpoints that shift destructive inflammation toward protective immune regulation remain poorly defined. Here, using the pristane-induced PH model, we identify interleukin-2-inducible T cell kinase (ITK) as a critical regulator of autoimmune lung injury. ITK deficiency (Itk⁻/⁻) conferred near-complete protection from PH and associated multiorgan injury, accompanied by reduced proinflammatory monocytes and neutrophils and increased Foxp3⁺ regulatory T cells (Tregs). Adoptive transfer of Itk⁻/⁻ Tregs protected wild-type recipients from PH and suppressed systemic proinflammatory cytokines, identifying Tregs as key mediators of tissue protection. Mechanistically, lung Itk⁻/⁻Tregs exhibited increased amphiregulin, a mediator of tissue repair. Transcriptomic profiling further showed that ITK loss reprogrammed Tregs toward a metabolically and functionally enhanced state, with enrichment of oxidative phosphorylation, mTORC1 and STAT5 signaling, and tissue-repair programs. These findings establish ITK as a regulator of the balance between pulmonary injury and reparative immunity and support targeting the ITK axis in severe inflammatory lung disease.

**Highlights:**

- ITK deficiency protects against pristane-induced pulmonary hemorrhage (PH)
- Loss of ITK expands “super-fit” canonical and non-canonical Tregs
- ITK-deficient Tregs from lungs exhibit enriched Amphiregulin production
- ITK-deficient Tregs exhibit enriched mTORC1, OXPHOS, and IL-10 production
- Transfer of ITK-deficient Tregs rescues established PH and reverses proteinuria
- ITK uncouples pathogenic inflammation from reparative tissue immunity

## Introduction

Pulmonary hemorrhage (PH) is a life-threatening manifestation of systemic autoimmunity, with reported mortality rates ranging from 40% to 90%. It can occur in autoimmune diseases such as anti-glomerular basement membrane disease (Goodpasture syndrome), systemic lupus erythematosus (SLE), and antiphospholipid syndrome, in which immune-mediated disruption of the alveolar–capillary interface causes rapid and potentially fatal pulmonary bleeding^1–3^. Current treatment generally relies on high-dose corticosteroids combined with systemic immunosuppressive agents, including cyclophosphamide and rituximab, with antifibrinolytic therapy used as an adjunct in selected cases to promote hemostasis^3,4^. Despite these aggressive interventions, clinical outcomes remain poor, underscoring a major unmet need to distinguish the immune mechanisms that drive organ-specific tissue injury from those that promote protection, repair, and recovery^5–8^.

Interleukin-2-inducible T cell kinase (ITK), a member of the Tec family of tyrosine kinases, is an important checkpoint in the regulation of T-cell responses^9–11^. In CD8⁺ T cells, ITK positively regulates T-cell receptor (TCR) signaling while restraining homeostatic proliferation, metabolic activity, and effector function^12^. Accordingly, CD8⁺ T cells from ITK-deficient mice acquire an innate memory-like phenotype characterized by increased expression of CD44 and CD122, together with the transcription factors Eomes and T-bet^11,13,14^. ITK also has important functions within the CD4⁺ T-cell compartment, where calcium-dependent ITK signaling regulates the balance between T helper 17 (Th17) and regulatory T-cell (Treg) differentiation^15^. In addition, ITK restrains IL-2-driven expansion of Foxp3⁺ Tregs^16^. Consistent with these functions, Itk⁻/⁻CD4⁺ T cells intrinsically generate a higher frequency of functional Foxp3⁺ Tregs than their wild-type counterparts^12^.

We previously demonstrated that disruption of ITK signaling uncouples graft-versus-host disease (GVHD) from graft-versus-leukemia (GVL) activity, supporting the concept that ITK differentially regulates pathogenic and protective T-cell responses^17–19^. We further showed that ITK deficiency promotes the development of highly suppressive Tregs with enhanced functional capacity^20^. However, whether ITK-dependent Treg reprogramming can protect the alveolar–capillary barrier during acute autoimmune lung injury remains unknown.

ITK functions as a proinflammatory amplifier in autoimmunity by strengthening T-cell receptor (TCR) signaling, promoting the differentiation of pathogenic T helper 17 (Th17) and T follicular helper (Tfh) cells, and restraining the development and suppressive function of Tregs^21,22^. Through these effects, ITK shifts the immune balance away from tolerance and toward persistent inflammation and tissue injury. Consistent with this role, enhanced ITK signaling in CD4⁺ T cells have been implicated in several autoimmune diseases, including rheumatoid arthritis and multiple sclerosis, positioning ITK as an important molecular checkpoint and a potential therapeutic target for restoring immune homeostasis^10,23^.

Despite these established functions, the contribution of ITK signaling to pristane-induced PH has not been investigated. Therefore, the central objective of this study was to define the cellular and molecular mechanisms through which ITK deficiency protects against severe autoimmune lung injury. Using Itk⁻/⁻ mice^17,19^, we show that loss of ITK profoundly remodels the T-cell compartment. Consistent with previous studies, both CD8⁺ and CD4⁺ T cells from Itk⁻/⁻ mice exhibited significant expansion of central memory and effector memory populations compared with wild-type controls.

To define the functional contribution of ITK to autoimmune lung injury, we challenged wild-type (WT) and Itk⁻/⁻ mice with pristane. We found that loss of ITK signaling established a dominant regulatory immune environment that protected mice from pulmonary hemorrhage and broadly attenuated systemic autoimmune injury^24^. This protection was characterized by reduced infiltration of proinflammatory immune cells and decreased circulating inflammatory cytokines, together with significant expansion of canonical CD4⁺CD25⁺Foxp3⁺ and noncanonical CD4⁺CD25⁻Foxp3⁺ Treg populations. Notably, Tregs from Itk⁻/⁻ mice exhibited increased expression of amphiregulin (AREG), an important mediator of tissue protection and repair ^25,26^.

Adoptive transfer of Itk⁻/⁻ Tregs protected WT recipients from PH and suppressed pathogenic inflammation, demonstrating their direct contribution to disease control. Transcriptomic profiling further revealed that Itk⁻/⁻ Tregs were reprogrammed toward a metabolically and functionally enhanced state. Together, these findings identify ITK as a previously unrecognized regulator of the balance between alveolar–capillary injury and reparative immunity and highlight the ITK pathway as a potential therapeutic target in severe inflammatory lung disease.

## Results

### ITK Deficiency Alters the Distribution of Naïve and Memory T-Cell Subsets

ITK is a Tec-family tyrosine kinase required for optimal T-cell receptor (TCR) signaling and plays an important role in T-cell development, activation, and effector function^27^.

To determine how ITK deficiency alters the T-cell compartment before autoimmune challenge, we analyzed splenic T-cell subsets from wild-type (WT) and Itk⁻/⁻ mice on a C57BL/6 background^17,18^. Because the differentiation state of αβ T cells strongly influences their effector potential and contribution to disease^28–31^, freshly isolated splenic CD3⁺ T cells were profiled by flow cytometry. CD3⁺ T cells were first separated into CD4⁺ and CD8⁺ populations and subsequently classified as naïve (CD44^lo^ CD62L^hi^), central memory (CM; CD44^hi^CD62L^hi^), or effector memory (EM; CD44^hi^CD62L^lo^) subsets^20,32^

Within the CD8⁺ T-cell compartment, Itk⁻/⁻ mice exhibited a significantly increased frequency of CM cells compared with WT controls, whereas the frequency of EM cells did not differ significantly between the two groups (Fig. 1A–D). Differences were more pronounced within the CD4⁺ T-cell compartment, in which Itk⁻/⁻mice displayed significant increases in both CM and EM populations, accompanied by a corresponding reduction in the naïve population (Fig. 1E–H; gating strategy shown in Figure 1—figure supplement 1). Together, these findings demonstrate that ITK deficiency alters the steady-state distribution of naïve and memory T-cell subsets, consistent with previous observations from our laboratory and others^12,14,16,33,34^.

**Figure 1.**
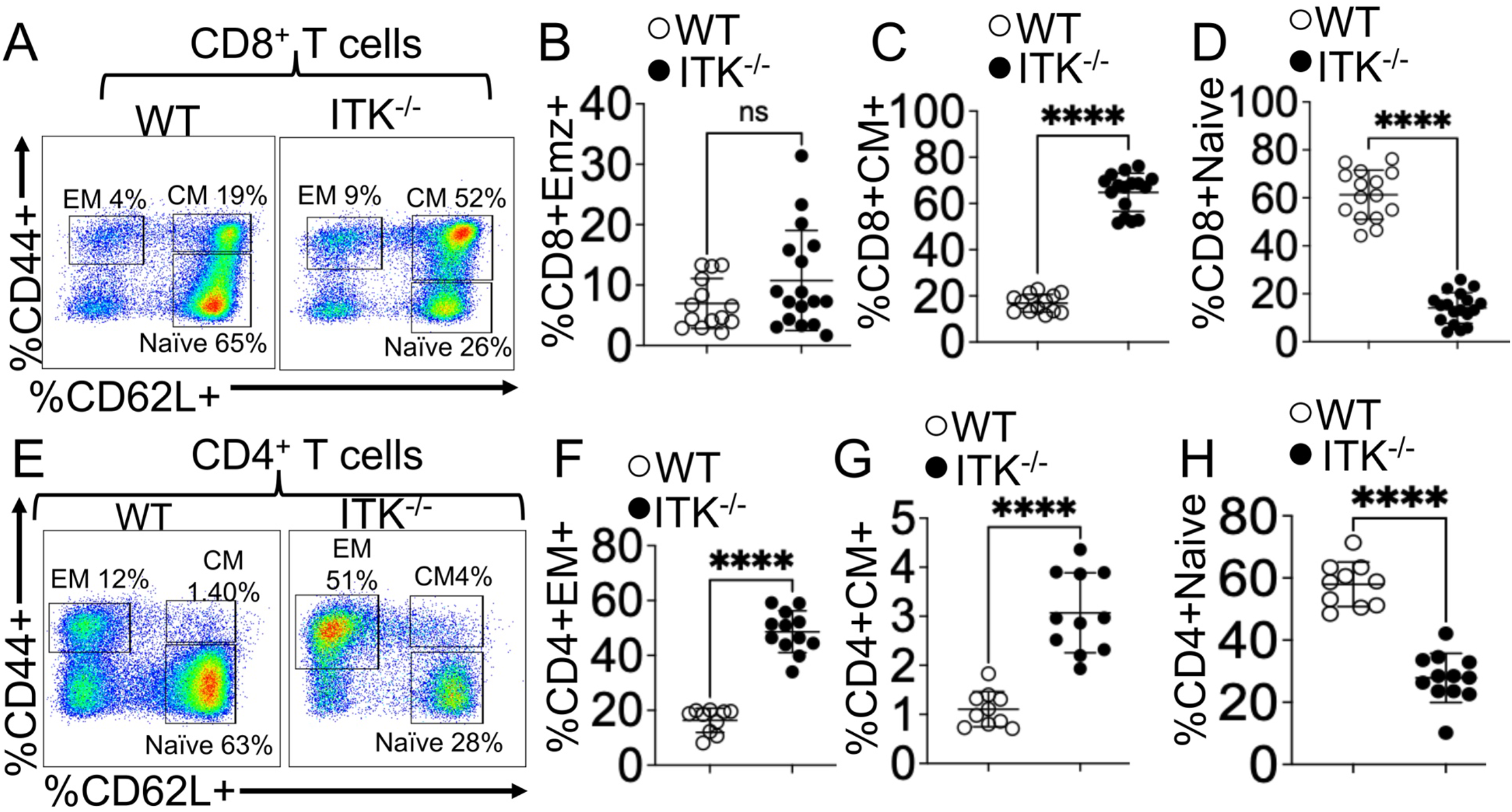
ITK deficiency promotes a predominantly memory-like T cell phenotype. (A–D) Analysis of the CD8⁺ T cell compartment in WT and Itk⁻/⁻ mice. Freshly isolated splenocytes were gated on CD3⁺ lymphocytes and further subdivided into CD8⁺ and CD4⁺ T cell populations. (A) Representative flow cytometry plots showing CD44 and CD62L expression within CD8⁺ T cells. (B–D) Quantification of effector memory (EM; CD44⁺CD62L⁻), central memory (CM; CD44⁺CD62L⁺), and naïve (CD44⁻CD62L⁺) CD8⁺ T cell subsets. (E–H) Analysis of the CD4⁺ T cell compartment. (E) Representative flow cytometry plots showing CD44 and CD62L expression within CD4⁺ T cells. (F–H) Quantification of EM, CM, and naïve CD4⁺ T cell frequencies. Data are presented as mean ± SEM (*n* = 10–15 mice per group). Results are representative of 6 independent experiments. Statistical significance was determined by an unpaired two-tailed Student’s *t*-test. \*\*\*\**p* ≤ 0.0001; ns, not significant (*p* > 0.05).

### ITK Deficiency Increases the Expression of CD44, CD122, T-bet, and Eomes in T Cells

We and others have previously shown that attenuation of T-cell receptor (TCR) signaling can alter T-cell activation and differentiation without necessarily promoting pathogenic alloimmune responses^17,18,20,35,36^. Consistent with these findings, flow cytometric analysis demonstrated significantly increased CD44 expression in splenic CD3⁺CD8⁺ T cells from Itk⁻/⁻mice compared with wild-type (WT) controls (Fig. 2A–B).

**Figure 2.**
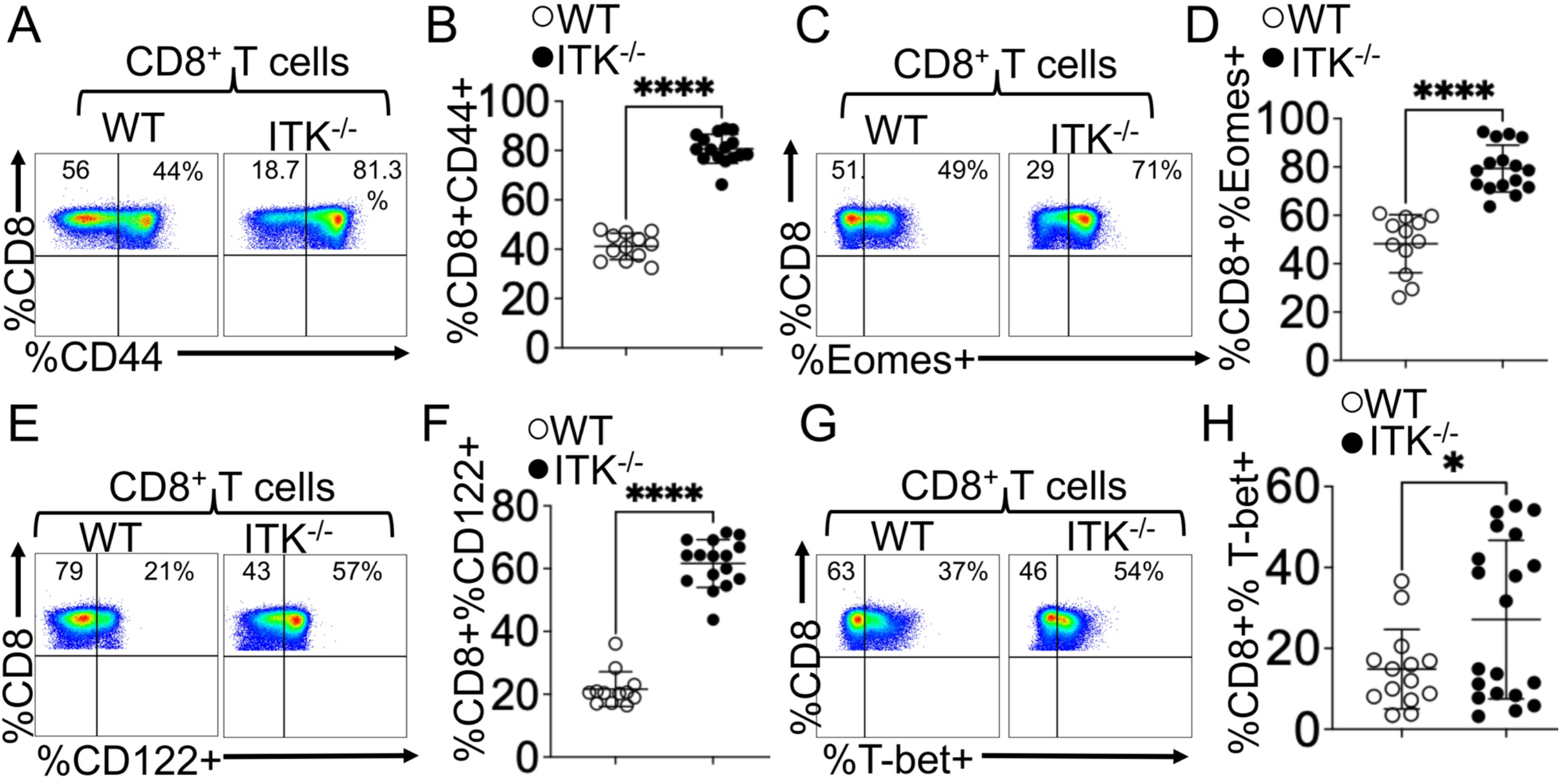
ITK deficiency enhances expression of memory-associated activating markers and transcriptional regulators. (A and B) Surface expression of CD44 on splenic T cells. Representative flow cytometry plots (A) and quantification of frequencies (B) in WT and Itk⁻/⁻ mice. (C and D) Expression of the T-box transcription factor Eomes. Representative flow cytometry plots (C) and quantification of total Eomes⁺ T cells (D). (E and F) Surface expression of CD122 (IL-2Rβ). Representative flow cytometry plots (E) and quantification of CD122 frequencies on T cells (F) from WT and Itk⁻/⁻mice. (G and H) Intracellular expression of the transcription factor T-bet. Representative flow cytometry plots (G) and quantification of T-bet frequencies within the indicated population (H). Data are presented as mean ± SEM (*n* = 15–20 mice per group, as indicated in the panels). Results are representative of at least 3 independent experiments. Statistical significance was determined by an unpaired two-tailed Student’s *t*-test. \**p* ≤ 0.05; \*\*\*\**p* ≤ 0.0001.

We next examined transcription factors and surface receptors associated with T-cell differentiation and cytokine responsiveness. The T-box transcription factors T-bet and Eomes regulate multiple aspects of effector and memory T-cell development and are associated with the expression of CD122 (IL-2Rβ), a shared component of the IL-2 and IL-15 receptor complexes^37,38^. CD8⁺ T cells from Itk⁻/⁻ mice exhibited significantly increased expression of T-bet, Eomes, and CD122 compared with WT controls (Fig. 2C–H). A similar pattern was observed in the CD4⁺ T-cell compartment, in which Itk⁻/⁻T cells displayed increased expression of CD44, Eomes, CD122, and T-bet (Figure 2—figure supplement 1A–H). Together, these findings demonstrate that ITK deficiency increases the expression of differentiation-associated transcription factors and surface markers in both CD8⁺ and CD4⁺ T-cell populations. These results further indicate that reduced ITK-dependent TCR signaling does not uniformly diminish the expression of molecules associated with T-cell differentiation and responsiveness to IL-2 and IL-15.

### ITK Deficiency Protects Against Pristane-Induced Pulmonary Hemorrhage and Systemic Pathology

We and others have previously shown that disruption of ITK signaling attenuates pathogenic immune responses in multiple models of alloimmunity ^17,18,20,35,36^. To investigate the role of ITK in autoimmune acute lung injury, we used the pristane-induced PH model, which reproduces key features of immune-mediated alveolar–capillary injury observed in systemic autoimmune diseases^1-3^. WT and Itk⁻/⁻ mice received a single intraperitoneal injection of pristane (0.5 mL) and were evaluated 14 days later, as established in previous studies^24,28,39,40^ (Fig. 3A). Neither group exhibited significant weight loss during this acute period. However, gross and histopathologic analyses at day 14 revealed marked differences in pulmonary injury. Lungs from WT mice exhibited extensive diffuse hemorrhage, intra-alveolar erythrocyte accumulation, and disruption of normal lung architecture. In contrast, Itk⁻/⁻ mice were protected from PH and maintained preserved alveolar architecture with minimal evidence of hemorrhage or tissue injury (Fig. 3B–E).

**Figure 3.**
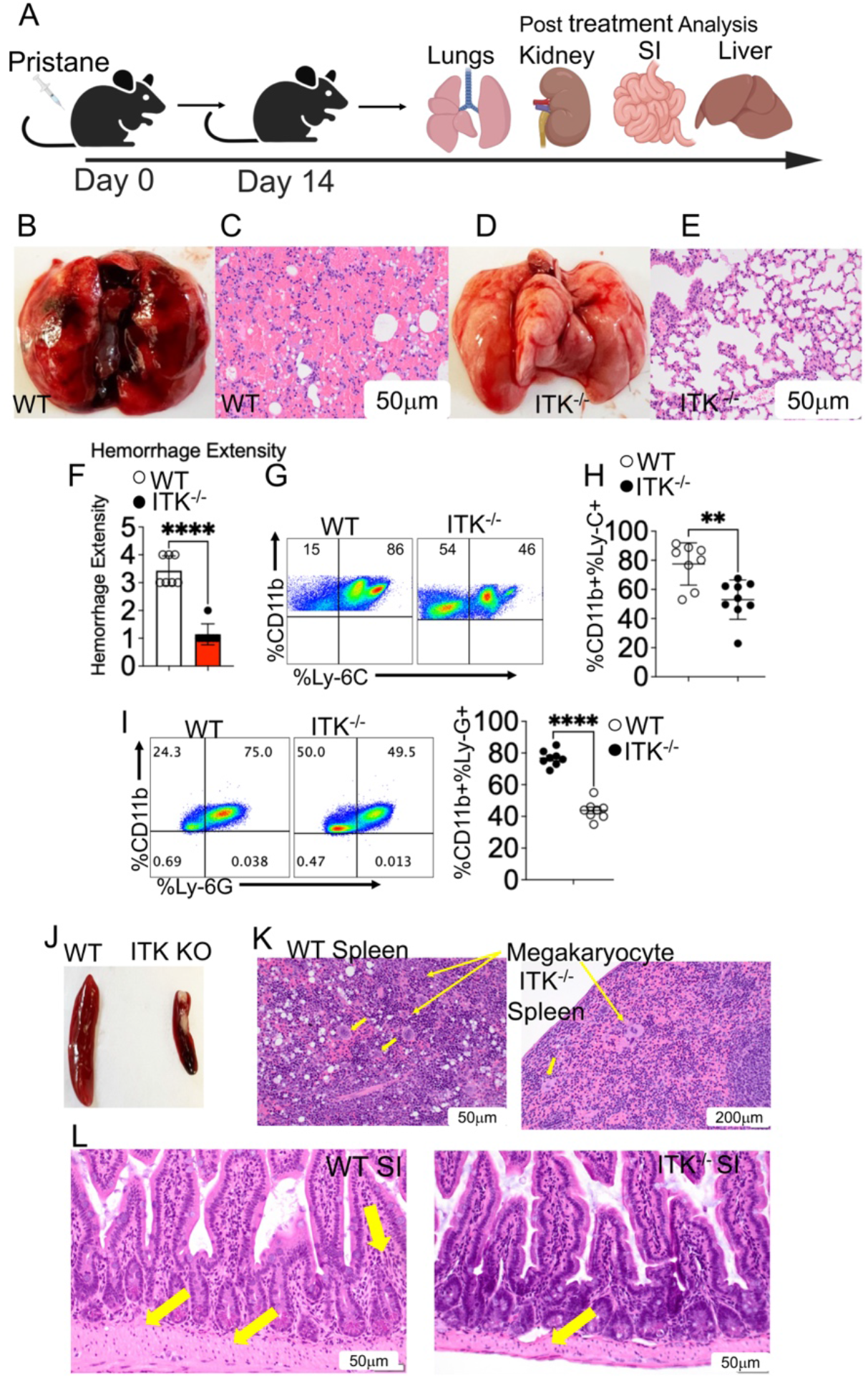
TK deficiency confers protection against pristane-induced PH and systemic pathology. (A) Experimental schematic: WT and Itk⁻/⁻ mice received a single i.p. injection of 0.5 mL pristane to induce pulmonary hemorrhage (PH) and were euthanized on day 14 for tissue analysis. (B–E) Assessment of lung pathology. Representative gross morphology (B, D) and H&E-stained lung sections (C, E) from WT and Itk⁻/⁻ mice after pristane injection. WT mice exhibit severe pulmonary hemorrhage and loss of architectural integrity, whereas lungs from Itk⁻/⁻ mice remain largely free of blood and retain preserved alveolar structures. (F) Quantification of blinded histopathologic scoring. Lung injury was graded by a board-certified pathologist blinded to genotype and treatment. (G and H) Analysis of lung-infiltrating proinflammatory monocytes. Representative flow cytometry plots (G) and quantification (H) of inflammatory monocytes, gated as CD11b⁺Ly6C⁺ within live singlets. (I) Representative flow cytometry plots and quantification of neutrophils (CD11b⁺Ly6G⁺) within live singlets from WT and Itk⁻/⁻ mice after pristane injection. (J and K) Evaluation of splenic pathology. Representative gross images (J) and H&E-stained sections (K) of spleens from WT and Itk⁻/⁻ mice after pristane injection. (L) Histopathological analysis of systemic inflammation. Representative H&E-stained sections of the small intestine from WT and Itk⁻/⁻mice. Data are presented as mean ± SEM (*n* = 10 mice per group). Results are representative of 3 independent experiments. Statistical significance was determined by an unpaired two-tailed Student’s *t*-test, except for panel F, which was analyzed using the Mann-Whitney *U* test. *p* ≤ 0.01; ** P ≤ 0.01, \*\*\*\**p* ≤ 0.0001.

To exclude the possibility that these differences reflected preexisting abnormalities, we examined lungs from untreated WT and Itk⁻/⁻ mice. Both groups exhibited normal pulmonary architecture, indicating that the protection observed in Itk⁻/⁻ mice emerged following pristane challenge rather than from baseline histologic differences (Figure 3—figure supplement 1A–D). We next assessed longer-term disease progression. All pristane-treated WT mice died within 20 days, whereas Itk⁻/⁻ mice exhibited significantly prolonged survival (Figure 3—figure supplement 2). Together, these findings demonstrate that ITK deficiency protects against acute pristane-induced PH and substantially improves survival.

Because the progression of PH is driven in part by the recruitment of pathogenic innate immune cells^41–43^, we next quantified inflammatory monocyte and neutrophil accumulation in the lungs. WT mice exhibited a marked influx of CD11b⁺Ly6C⁺ inflammatory monocytes following pristane challenge, whereas this population was significantly reduced in Itk⁻/⁻ mice (Fig. 3F–H). Because neutrophils are early responders to inflammatory tissue injury^43,44^, we also assessed their accumulation in the lungs at day 14 after pristane administration. CD11b⁺Ly6G⁺ neutrophils were significantly reduced in Itk⁻/⁻ mice compared with WT controls (Fig. 3I). Together, these findings demonstrate that protection from pristane-induced PH in Itk⁻/⁻ mice is associated with reduced pulmonary accumulation of proinflammatory innate immune cells.

Beyond the lung, ITK deficiency also attenuated systemic manifestations of pristane-induced injury^45,46^. Following pristane challenge, WT mice developed pronounced splenomegaly accompanied by extensive accumulation of megakaryocytes, whereas spleens from Itk⁻/⁻mice retained near-normal size and histologic architecture (Fig. 3J–L; Figure 3—figure supplement 3E–F). Histologic examination of spleens from naïve WT and Itk⁻/⁻ mice revealed no baseline abnormalities, indicating that these differences emerged following pristane exposure (Figure 3—figure supplement 3A–C).

We next examined the small intestine. Naïve WT and Itk⁻/⁻mice exhibited normal intestinal architecture without evidence of preexisting tissue injury (Figure 3—figure supplement 4A–B). Following pristane administration, WT mice developed substantial small-intestinal pathology characterized by epithelial hyperplasia and focal acute inflammation. These changes were markedly attenuated in Itk⁻/⁻mice (Fig. 3M; Figure 3—figure supplement 4C–D).

Together, these findings demonstrate that ITK deficiency provides multiorgan protection against pristane-induced systemic inflammatory injury. Having established this protective phenotype, we next examined whether loss of ITK also reduced systemic proinflammatory cytokine production.

### ITK Deficiency Attenuates Proteinuria and Shifts the Systemic Cytokine Balance Toward Resolution

Proteinuria is a clinical hallmark of pulmonary-renal syndrome and frequently accompanies PH in systemic autoimmune disease, serving as an important indicator of glomerular involvement^47^. To determine whether the protection observed in Itk⁻/⁻ mice extended to the kidney, we quantified urinary protein levels at baseline (day 0) and at the study endpoint (day 14). Whereas WT mice developed a significant increase in proteinuria following pristane challenge, ITK⁻/⁻ mice showed a markedly attenuated response, indicating that loss of ITK signaling protects against PH-associated renal dysfunction (Fig. 4A-C). Histologic analysis at this acute 14-day time point did not reveal overt structural kidney injury, although WT mice displayed focal inflammatory cell infiltration that was absent in ITK⁻/⁻ mice (Supplementary Fig. 5). These findings suggest that although glomerular injury remains at an early stage in this acute model, the ITK axis already regulates the inflammatory processes that precede overt renal damage.

**Figure 4.**
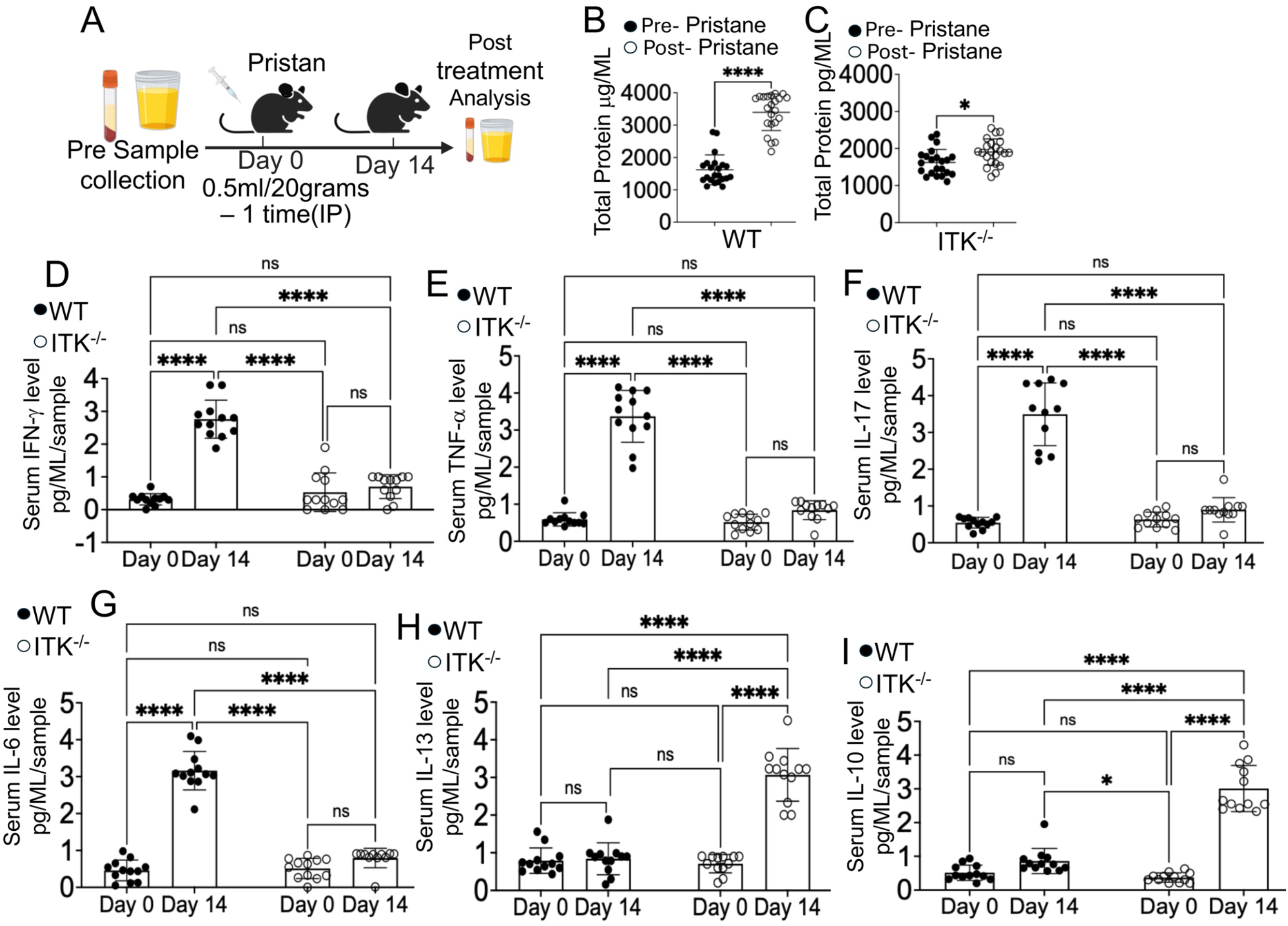
ITK deficiency attenuates proteinuria and shifts the systemic cytokine balance toward resolution. (A) Experimental schematic: WT and Itk⁻/⁻mice were challenged with pristane. Urine and blood were collected at baseline (day 0) and at the day 14 endpoint to assess systemic pathology. (B and C) Assessment of renal function by proteinuria. Total urinary protein was quantified by BCA assay in WT (B) and Itk⁻/⁻ mice (C). Itk⁻/⁻ mice show significant protection against the glomerular leakage observed in WT controls following pristane injection. (D–I) Multiplex serum cytokine profiling. Serum levels of proinflammatory mediators (IFN-γ, TNF-α, IL-17, and IL-6) and pro-resolving cytokines (IL-13 and IL-10) were quantified by bead-based immunoassay. Itk⁻/⁻mice exhibited marked suppression of proinflammatory cytokines together with preservation of regulatory cytokines. Data are presented as mean ± SEM (*n* = 15–25 mice per group). Results are representative of 3 independent experiments. Statistical significance was determined by a two-tailed Student’s *t*-test for single comparisons or a two-way ANOVA with Tukey’s *post hoc* correction for time-course data. *p* < 0.01; \*\*\**p* < 0.001; \*\*\*\**p* < 0.0001, ** P ≤ 0.01

Because PH is associated with a robust systemic inflammatory response, we next quantified circulating cytokines to determine how the ITK axis influences the balance between pathogenic and reparative immunity. We measured key mediators implicated in PH pathogenesis, including IFN-γ and TNF-α, which promote procoagulant activity and tissue factor expression^48^, as well as IL-6 and IL-17, which drive neutrophil recruitment and vascular permeability^49–53^. Serum analysis by ELISA by (LEGENDplex/multiplex cytokine analysis), showed that ITK⁻/⁻mice had significantly lower levels of all four proinflammatory cytokines than WT controls (Fig. 4D-G). Conversely, because anti-inflammatory cytokines can restrain pathogenic immune responses^54,55^, we asked whether ITK deficiency also promotes a pro-resolving cytokine environment. Indeed, ITK⁻/⁻mice produced significantly higher levels of IL-10 and IL-13 during PH (Fig. 4H-I).

Together, these findings indicate that ITK deficiency does not simply impose broad immunosuppression, but instead redirects the systemic immune response toward regulation and resolution, further supporting ITK as a promising therapeutic target in PH and related autoimmune diseases.

### Itk⁻/⁻ Tregs Exhibit Enhanced AREG Expression and More Effectively Ameliorate PH

Regulatory T cells (Tregs) are key mediators of immune suppression and tissue repair, acting in part through anti-inflammatory cytokines such as IL-10 to restrain neutrophils and inflammatory monocytes^56,57^. We previously demonstrated that ITK deficiency increases the frequency of both canonical (CD4⁺CD25⁺Foxp3⁺) and non-canonical (CD4⁺CD25⁻Foxp3⁺) Tregs^19,58^. To determine whether these regulatory populations are similarly altered during autoimmune lung injury, we analyzed splenic Tregs from WT and Itk⁻/⁻mice following pristane challenge.

Itk⁻/⁻ mice exhibited significantly higher frequencies of both canonical and non-canonical Treg subsets compared with WT controls (Fig. 5A; Supplementary Fig. 7A–B). These findings are consistent with the established role of ITK in constraining the regulatory T-cell compartment and demonstrate that this altered Treg distribution is maintained during pristane-induced systemic inflammation. Notably, the increased representation of the non-canonical CD25⁻Foxp3⁺ population is consistent with our previous studies demonstrating enrichment and suppressive activity of this population in the setting of ITK deficiency^12,59,60^. We next examined whether this altered regulatory phenotype was also present in the lung.

**Figure 5.**
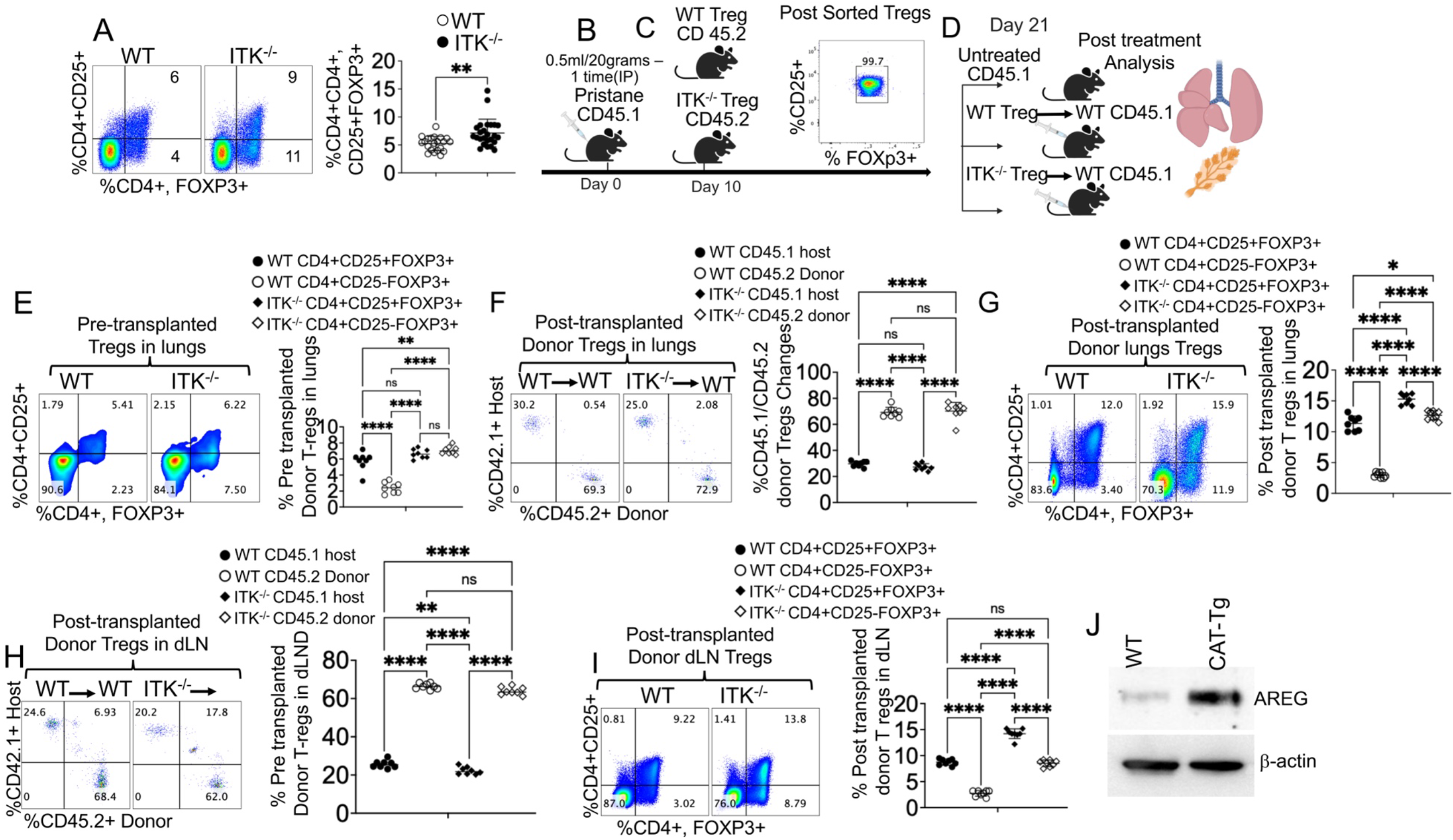
Adoptively transferred Itk⁻/⁻Tregs exhibit enhanced AREG expression and more effectively ameliorate PH. (A) Representative flow cytometry plots and quantitative analysis of splenic CD4⁺Foxp3⁺ Tregs in WT and Itk⁻/⁻mice. (B–D) Experimental schematic for Treg adoptive transfer. (B) WT mice (CD45.1⁺) received a single i.p. injection of 0.5 mL pristane. (C) On day 10, CD4⁺Foxp3⁺ Tregs were FACS-sorted from WT or Itk⁻/⁻ donor mice (CD45.2⁺), with post-sort purity confirmed as CD25⁺Foxp3⁺. These cells were injected intravenously (i.v.) into the pristane-treated CD45.1⁺ recipients as a therapeutic intervention. (D) On day 21, CD45.1⁺ host mice were euthanized for examination of the lungs and draining lymph nodes (dLNs). (E) Representative flow cytometry plots and quantitative analysis of lung CD4⁺Foxp3⁺ Tregs in WT and Itk⁻/⁻ mice at baseline (without pristane injection). (F and G) Representative flow cytometry plots and quantitative analysis of total donor CD45.2⁺ cells (F) and donor-derived Tregs (G) engrafted in the lungs of CD45.1⁺ recipient mice. (H and I) Representative flow cytometry plots and quantitative analysis of total donor CD45.2⁺ cells (H) and donor-derived Tregs (I) in the dLNs of CD45.1⁺ recipient mice. (J) Immunofluorescence microscopy of lung sections confirming the presence of donor Tregs in inflamed lung tissue. Sections were stained for CD4, the donor congenic marker (CD45.2), and Foxp3, with merged images demonstrating colocalization; nuclei were counterstained with DAPI. (K) CD25⁺Foxp3⁺ Tregs were FACS-sorted from lung tissue and analyzed by immunoblot for amphiregulin (AREG), with β-actin as a loading control. Data are presented as mean ± SEM; sample sizes (*n* = 15–25 mice per group) are indicated in the panels. Each experiment was repeated 3 times. Statistical significance was determined using a two-tailed Student’s *t*-test, one-way ANOVA, or two-way ANOVA as appropriate; *p* < 0.01, \*\*\**p* < 0.001, * P ≤ 0.05 \*\*\*\**p* < 0.0001, ** P ≤ 0.01.

Analysis of lung tissue under homeostatic conditions revealed an increased frequency of Tregs in *Itk⁻/⁻*mice compared with WT controls, with the most pronounced increase observed in the non-canonical CD25⁻Foxp3⁺ subset (Fig. 5E). From approximately 1 × 10⁷ total viable lung cells recovered per mouse, approximately 8 × 10⁵ CD3⁺CD4⁺ T cells were identified in both WT and Itk⁻/⁻mice, indicating comparable overall CD4⁺ T-cell abundance between genotypes. Absolute Treg numbers were calculated from the total CD3⁺CD4⁺ T-cell recovery and the corresponding Treg frequencies determined by flow cytometry. In WT lungs, canonical CD25⁺Foxp3⁺ Tregs comprised approximately 6% of CD4⁺ T cells, corresponding to approximately 4.8 × 10⁴ cells, whereas non-canonical CD25⁻Foxp3⁺ Tregs comprised approximately 2.3%, corresponding to approximately 1.8 × 10⁴ cells. In Itk⁻/⁻lungs, canonical Tregs comprised approximately 6–7%, corresponding to approximately 4.8–5.6 × 10⁴ cells, whereas non-canonical Tregs increased to approximately 8%, corresponding to approximately 6.4 × 10⁴ cells. Thus, the increased Treg frequencies in Itk⁻/⁻ mice were not attributable to reduced overall CD4⁺ T-cell abundance and were accompanied by a marked increase in the absolute number of non-canonical CD25⁻Foxp3⁺ Tregs.

To determine whether Itk⁻/⁻ Tregs possess enhanced protective activity *in vivo*, we performed adoptive-transfer experiments in WT recipients following initiation of pristane-induced disease. WT CD45.1 congenic mice were challenged with pristane and monitored for 10 days. On day 10, mice were assigned to three groups: a pristane-challenged control group receiving no Tregs, a group receiving FACS-purified WT CD45.2⁺ Tregs (CD3⁺CD4⁺CD25⁺Foxp3⁺), and a group receiving an equivalent number of Itk⁻/⁻ CD45.2⁺ Tregs (1× 10⁶ cells per mouse; Fig. 5B–D). Tregs were isolated from the spleens of **unchallenged** WT or Itk⁻/⁻ Foxp3-RFP reporter mice, allowing purification of viable CD25⁺Foxp3⁺ Tregs as previously described^19,24^. On day 21, recipient mice were euthanized, and the lungs and draining lymph nodes (dLN) were analyzed (Fig. 5D).

We next examined donor-derived Tregs by distinguishing CD45.1⁺ recipient cells from CD45.2⁺ donor cells. At the experimental endpoint, a significantly higher frequency of donor-derived Itk⁻/⁻ Tregs was detected in the lungs compared with WT donor Tregs (Fig. 5F). To determine the phenotypic composition of the CD45.2⁺ donor population after transfer, we assessed Foxp3 and CD25 expression. Both canonical CD25⁺Foxp3⁺ and non-canonical CD25⁻Foxp3⁺ donor-derived populations were detectable, with higher frequencies observed among cells derived from Itk⁻/⁻ donors compared with WT donors (Fig. 5G).

Consistent with the lung findings, we examined donor-derived Tregs in the dLN of WT CD45.1 recipients^18,19^. A significantly higher frequency of CD45.2⁺ donor-derived Itk*⁻/⁻* Tregs was detected in the dLN compared with WT donor Tregs (Fig. 5H). Within the donor-derived CD45.2⁺CD3⁺CD4⁺Foxp3⁺ population, both canonical CD25⁺ and non-canonical CD25⁻ subsets were identified, with greater representation among Itk⁻/⁻ donor cells (Fig. 5I).

To further characterize the tissue-repair-associated phenotype of Itk⁻/⁻ Tregs, we evaluated Amphiregulin (AREG), an epidermal growth factor receptor ligand implicated in tissue repair independently of classical Treg suppressive functions^23,61–63^. CD25⁺Foxp3⁺ Tregs were FACS-sorted from the lungs of WT and Itk⁻/⁻ Foxp3-RFP reporter mice, and AREG protein expression was assessed by immunoblotting. Tregs isolated from Itk⁻/⁻ lungs expressed substantially higher levels of AREG than their WT counterparts (Fig. 5J), consistent with an enhanced tissue-repair-associated phenotype.

Collectively, these findings demonstrate that Itk⁻/⁻ Tregs are more prominently represented in disease-relevant tissues following transfer and exhibit increased AREG expression, providing additional phenotypic features associated with their enhanced protective activity during PH.

### Adoptively Transferred Itk⁻/⁻ Tregs Protect Mice from PH, Reduce Proinflammatory Cytokine Levels, and Increase Anti-inflammatory Cytokine Levels

Tregs suppress autoimmune disease primarily by secreting inhibitory cytokines. These specialized signaling proteins actively dampen harmful immune responses, maintain self-tolerance, and protect healthy tissues from autoimmune destruction^61,62^. In addition to their suppressive function, Tregs contribute directly to the restoration of the alveolar-capillary barrier following inflammatory injury^56,63^.

To determine whether the adoptive transfer of Tregs from WT or Itk⁻/⁻ mice reduced proinflammatory cytokines or increased anti-inflammatory cytokines, we repeated the *in vivo* transfer experiments described above (Fig. 5). WT (CD45.1) congenic mice were injected intraperitoneally with pristane (0.5 mL) to induce disease. On day 10, following the establishment of PH, CD25⁺Foxp3⁺ Tregs were FACS-sorted from the spleen of WT (CD45.2) and Itk⁻/⁻ (CD45.2) donor mice utilizing Foxp3-RFP reporter expression, and were subsequently transferred into the affected recipients (Fig. 6A-D).

**Figure 6.**
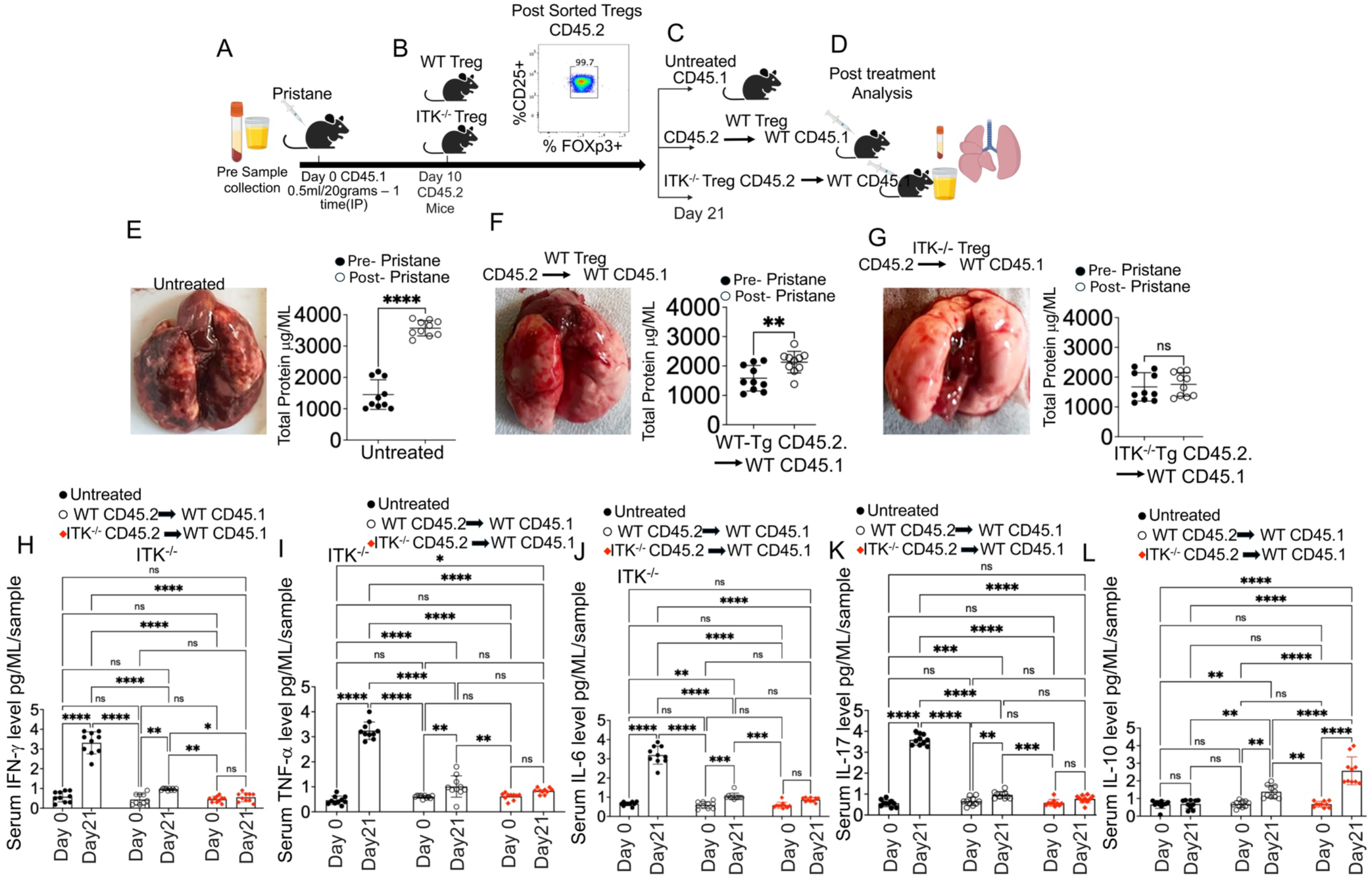
Adoptively transferred Itk⁻/⁻ Tregs protect mice from PH, reduce proinflammatory cytokine levels, and increase anti-inflammatory cytokine levels. (A–D) Experimental schematic for the therapeutic adoptive transfer model. (A) Baseline serum and urine were collected from WT host mice (CD45.1⁺), followed by a single i.p. injection of 0.5 mL pristane. (B and C) On day 10, CD4⁺Foxp3⁺ Tregs were FACS-sorted from WT or Itk⁻/⁻ donor mice (CD45.2⁺), confirmed positive for CD25 and Foxp3, and injected i.v. into the pristane-treated CD45.1⁺ recipients. A control group of pristane-challenged CD45.1⁺ mice was left untreated. (D) On day 21, host mice were euthanized and analyzed for urinary protein and gross anatomical damage to the lungs. (E–G) Evaluation of therapeutic efficacy in lung and kidney injury. Representative gross lung images and longitudinal quantification of urinary protein (day 0 versus day 21) are shown for (E) untreated pristane-challenged mice, (F) mice treated with WT Tregs, and (G) mice treated with Itk^-/-^ Tregs. (H–L) Suppression of systemic inflammatory responses. Serum levels of proinflammatory cytokines (IFN-γ, TNF-α, IL-6, and IL-17) and the pro-resolving mediator IL-10 were quantified at baseline and day 21. Itk⁻/⁻ Tregs promoted a high-IL-10 regulatory environment while effectively suppressing the proinflammatory cytokine response. Data are presented as mean ± SEM (*n* = 15–25 mice per group). Results are representative of 3 independent experiments. Statistical significance was determined by a two-tailed Student’s *t*-test or a two-way ANOVA with Tukey’s *post hoc* correction. ns, *p* > 0.05; \**p* ≤ 0.05; *p* ≤ 0.01; \*\*\*\**p* ≤ 0.001; ** P ≤ 0.01, \*\*\*\*\**p* ≤ 0.0001.

By day 21 post-pristane injection, untreated WT mice developed severe gross pulmonary pathology alongside a significant increase in proteinuria relative to their baseline (day 0) levels (Fig. 6E). Recipients of WT Tregs exhibited only partial protection; while they showed a moderate reduction in proteinuria compared with untreated controls, their protein levels remained significantly elevated above baseline (Fig. 6F). In striking contrast, WT mice treated with Itk⁻/⁻ Tregs were fully protected from PH, with their proteinuria maintained at pre-challenge baseline levels (Fig. 6G). Together, these findings demonstrate that Itk⁻/⁻ Tregs possess enhanced suppressive and tissue-protective capacity *in vivo*, further supporting the modulation of the ITK axis as a promising therapeutic strategy for PH and severe autoimmune inflammatory injury.

Consistent with the reduced cytokine levels observed in Itk⁻/⁻ mice (Fig. 4), we next asked whether the adoptive transfer of Itk⁻/⁻ Tregs could similarly reshape the systemic cytokine milieu in WT recipients. Serum ELISA analysis confirmed that pristane-challenged WT mice developed marked increases in IFN-γ, TNF-α, IL-6, and IL-17 (Fig. 6H-K). Although the transfer of WT Tregs produced a modest reduction in these proinflammatory cytokines, recipients of Itk⁻/⁻ Tregs exhibited significantly greater suppression of this inflammatory response. Notably, WT mice treated with Itk⁻/⁻Tregs also showed a significant increase in serum IL-10, a key mediator of immune regulation and resolution, exceeding the levels observed in both untreated and WT Treg-treated groups (Fig. 6L).

These findings indicate that Itk⁻/⁻ Tregs not only suppress pathogenic Th1- and Th17-associated inflammatory responses more effectively than WT Tregs, but also actively promote an anti-inflammatory systemic environment. To determine whether Itk⁻/⁻ Tregs provide better long-term disease resolution, the recipient mice were monitored for survival. Our data confirmed that WT mice transplanted with Itk⁻/⁻ Tregs exhibited significantly prolonged survival compared with recipients of WT Tregs. Collectively, these results demonstrate that Itk⁻/⁻ Tregs confer enhanced protection against PH by shifting the cytokine balance from a proinflammatory toward a pro-resolving state.

### ITK Deficiency Reprograms the Treg Transcriptome Toward a Metabolically and Functionally Enhanced State

Having established that Itk⁻/⁻ Tregs confer enhanced protection against PH, we next sought to define the global transcriptomic programs underlying this phenotype. We performed RNA sequencing (RNA-seq) on FACS-purified canonical (CD3⁺CD4⁺CD25⁺Foxp3⁺) Tregs from WT and Itk⁻/⁻mice, as well as non-canonical (CD3⁺CD4⁺CD25⁻Foxp3⁺) Tregs from Itk⁻/⁻mice. Notably, the non-canonical subset was selectively expanded in Itk⁻/⁻ mice, precluding direct comparison with an equivalent WT population. Hierarchical clustering of normalized gene expression profiles revealed distinct transcriptomic segregation, with samples partitioning into two principal modules: Module 1, consisting of genes downregulated in Itk⁻/⁻subsets, and Module 2, comprising genes upregulated in Itk⁻/⁻ Tregs (Fig. 7A-B).

**Figure 7.**
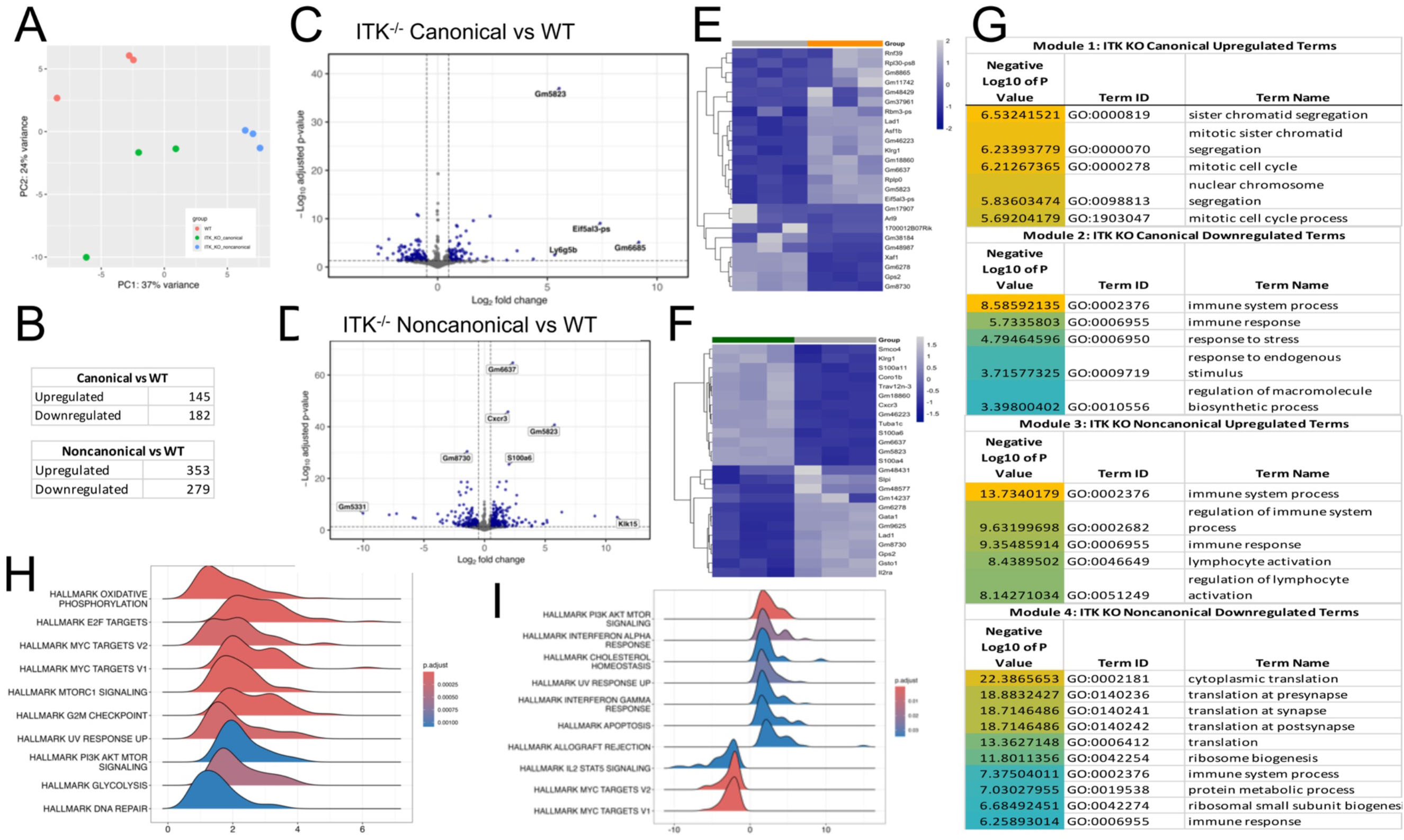
ITK deficiency reprograms the Treg transcriptome toward a metabolically and functionally altered state. (A) Global transcriptomic relationships. Principal component analysis (PCA) of transcriptomes from WT and Itk⁻/⁻ canonical (CD25⁺) and non-canonical (CD25⁻) Tregs. All replicates are shown (*n* = 3). (B) Table showing the number of up- or downregulated differentially expressed genes (DEGs) between groups. (C and D) Transcriptomic landscape of ITK-deficient Tregs. Volcano plots identify the top DEGs between Itk⁻/⁻and WT Tregs. Genes meeting the thresholds of FDR ≤ 0.05 and |log₂FC| ≥ 0.5 are highlighted, with a positive log₂FC indicating enrichment in Itk⁻/⁻ canonical (C) and non-canonical (D) Tregs. (E and F) Hierarchical clustering heatmaps showing row-scaled normalized expression of all significant DEGs in canonical CD4⁺CD25⁺Foxp3⁺ Tregs from WT and Itk⁻/⁻mice (*n* = 3 per group). A grey bar indicates WT, orange indicates the canonical group, and green indicates the non-canonical group. (G) Tables showing GO enrichment analysis of the DEGs from Itk⁻/⁻ canonical (CD25⁺) and non-canonical (CD25⁻) Tregs. Using the online tool gProfiler and the ordered g:GOSt query, we assessed which biological processes (BP) were linked to the genes in the different modules from Itk⁻/⁻and WT mice. In the table, *p*-values are color-coded from light blue (not significant) to orange (highest significance) (*n* = 3 mice per group). (H and I) Gene Set Enrichment Analysis (GSEA) of regulatory pathways in canonical (H) and non-canonical (I) samples. Ridge plots show the distribution of ranked *DESeq2* Wald statistic values for selected Hallmark pathways, including oxidative phosphorylation (OXPHOS), mTORC1 signaling, and MYC targets, which together reflect enhanced metabolic fitness in ITK-deficient regulatory subsets.

To identify the biological programs associated with the enhanced function of Itk⁻/⁻Tregs, we performed Gene Set Enrichment Analysis (GSEA) and hallmark pathway profiling. Dot plot and ridge plot analyses of normalized enrichment scores (NES) revealed significant enrichment of multiple pathways in both canonical and non-canonical Itk⁻/⁻ Tregs, including MYC targets, mTORC1 signaling, oxidative phosphorylation (OXPHOS), and glycolysis (Fig. 7C-H). Additional enrichment was observed in pathways related to DNA repair, E2F targets, and immune homeostasis, alongside the differential regulation of IFN-γ and NF-κB signaling. GSEA enrichment plots further supported the conclusion that the loss of ITK signaling promotes a metabolically active and functionally enhanced Treg state, characterized by increased cell cycle- and metabolism-associated programs (Supplementary Fig. 9A-K). Together, these data indicate that ITK deficiency does not simply expand Treg numbers, but instead reprograms the Treg transcriptome toward a state consistent with greater metabolic fitness and regulatory function. These findings provide a mechanistic framework for the potent tissue-protective activity of Itk*⁻/⁻* Tregs in PH.

### Gene-Level Analyses Reveal Distinct Transcriptional Programs in Canonical and Non-Canonical Itk⁻/⁻ Tregs

To address whether these pathway level differences reflect coordinated changes in identifiable genes, we examined the leading-edge genes of each enriched pathway (Figure 8). In canonical Itk*⁻/⁻* Tregs the oxidative phosphorylation signature was carried by ninety-two leading edge genes, including subunits of complex I (Ndufs8, Ndufb8), complex III (Uqcrc1, Cyc1), complex IV (Cox5b) and the mitochondrial ATP synthase (Atp5f1b, Atp5f1d, Atp5mc1, Atp5mc3). None of these genes reached statistical significance on its own, so this signature consists of small increases distributed across the respiratory chain rather than a change in any single enzyme, and it would not have been recovered by differential expression testing alone. Glycolysis and DNA repair were also enriched in canonical Itk*⁻/⁻* Tregs. The E2F and MYC target signatures, by contrast, were driven by genes with larger individual changes, including the replicative helicase components Mcm3, Mcm5 and Mcm7 together with Ccnb2, Birc5, Espl1, Asf1b and Mybl2. The four pathways drew on largely distinct gene sets, with 207 leading edge genes assigned to a single pathway and only three shared across three pathways, indicating that they are not restatements of a single signature.

**Figure.8.**
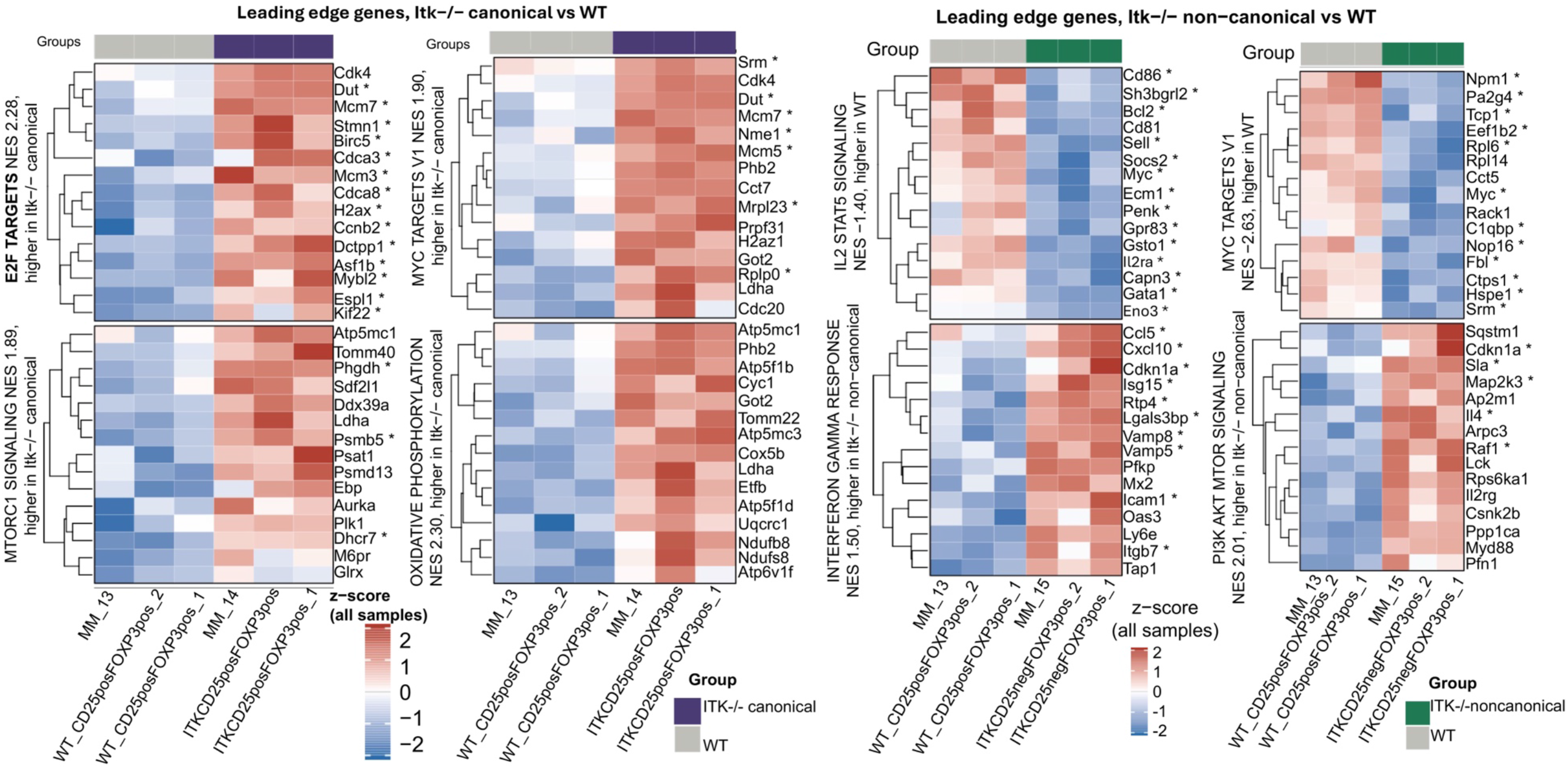
Gene-Level Analyses Reveal Distinct Transcriptional Programs in Canonical and Non-Canonical *Itk⁻/⁻*Tregs. (A, B) Leading edge genes of the Hallmark pathways shown in Figure 7, for Itk*⁻/⁻* canonical and Itk*⁻/⁻* non-canonical Tregs against WT canonical Tregs. Within each pathway the leading-edge genes, that is the core enrichment set returned by GSEA, were ranked by the absolute DESeq2 Wald statistic and the top fifteen are shown. Color is variance stabilized, batch corrected expression, z-scored across all nine samples, so the two panels are directly comparable. Asterisks mark genes that are also individually differentially expressed (FDR ≤ 0.05, |log2FC| ≥ 0.5). Row headings give the normalized enrichment score and the group in which each pathway is higher. (C, D, E) Genes annotated to the most significant Gene Ontology biological process terms for the differentially expressed genes in each comparison, tested against a background of the 16,723 expressed genes. Up to eight terms are shown per comparison and up to twelve genes per term, ranked by absolute fold change; color is the apeglm shrunken log2 fold change. No biological process term reached significance among the genes downregulated in canonical Itk−/− Tregs against this background.

Gene Ontology analysis of the genes upregulated in canonical Itk*⁻/⁻* Tregs returned terms centered on sister chromatid segregation, mitotic nuclear division and spindle attachment, driven by Espl1, Xrcc3, Spag5, Birc5, Ncaph, Mybl2, Cdca8 and Cdk1. No biological process term reached significance among the downregulated genes in this comparison.

Non-canonical Itk*⁻/⁻* Tregs showed a partly opposing profile. MYC targets and IL2/STAT5 signalling were enriched in wild type rather than in Itk*⁻/⁻* cells, the latter consistent with the CD25 negative phenotype used to define this subset, while PI3K/AKT/mTOR signalling and the interferon gamma response were enriched in Itk*⁻/⁻* cells. The upregulated genes were annotated to lymphocyte and alpha beta T cell activation, leukocyte cell–cell adhesion and chemokine-mediated signalling, with Ascl2 showing by far the largest change, followed by Il4, Cxcr3, Prdm1, Ccr2, Tigit, Icos, Cxcr5, Tbx21 and Runx3. The downregulated genes were dominated by ribosome biogenesis and cytoplasmic translation, including Npm1, Dkc1, Nop58 and numerous ribosomal protein genes. Gene significant hallmark, leading Edge gene among Itk*⁻/⁻* Non-canonical Tregs and Canonical, leading, ploted Canonical, GSVE per samples, GO Itk*⁻/⁻* Non-canonical Tregs and Canonical are listed in supplementary tables

## Discussion

Pulmonary hemorrhage (PH) remains a life-threatening clinical emergency in systemic autoimmune disease. With mortality rates frequently exceeding 50%, there is a pressing need to move beyond broad immunosuppression toward therapeutic strategies that promote immune resolution and tissue protection^64–68^. In this study, we identify ITK as a key molecular checkpoint that regulates the balance between destructive pulmonary inflammation and reparative immunity. Our findings demonstrate that ITK deficiency confers marked protection against PH and is associated with a broad remodeling of the T cell compartment toward a dominant regulatory state with features of enhanced metabolic fitness.

ITK has long been viewed as a signaling rheostat, whereby its absence or inhibition attenuates T cell receptor (TCR) signaling in ways that favor the development of memory CD8⁺ T cells and regulatory T cell lineages over pathogenic Th17 and Tfh populations^9,12,19,23,69,70^. Although inherited ITK deficiency in humans is associated with impaired IFN-γ immunity and defective viral control^71,72^, our findings suggest that in the setting of acute autoimmune injury, reduced ITK signaling does not result in immune failure, but instead supports a more protective regulatory response. In particular, we show that Itk⁻/⁻ Tregs are not only expanded in number, but are also sufficient to rescue established PH in wild-type recipients. This regulatory reprogramming may help explain why ITK-deficient T cells, which we previously showed retain anti-tumor activity without inducing graft-versus-host disease (GVHD) ^17^, are also capable of protecting the alveolar-capillary barrier during autoimmune injury.

Our study further shows that ITK deficiency reshapes the T cell compartment, characterized by the expansion of effector memory (EM) and central memory (CM) populations in both the CD4⁺ and CD8⁺ lineages. This memory-like shift is accompanied by increased expression of activation-associated markers and the T-box transcription factors Eomes and T-bet, which are known to support effector competence and durable memory differentiation^37,73^. Interestingly, although T-bet is typically induced downstream of TCR, IL-12, and mTOR signaling, and can promote Eomes expression through IL-2/STAT5-dependent pathways^34,37,74–76^, our findings suggest that ITK deficiency creates a distinct signaling context that favors the acquisition of an Eomes-rich, memory-like phenotype. While the role of ITK in T cell development is well established^11,13,77^, our results extend its significance to systemic autoimmunity and inflammatory lung injury. We propose that this preconfigured memory-like state may contribute to protection during acute injury by enabling more controlled and effective immune responses, thereby limiting the unchecked innate inflammation that characterizes pristane-induced PH.

Utilizing the pristane-induced murine model, we show that ITK deficiency confers marked protection against PH, as evidenced by the preservation of lung architecture and a striking reduction in pulmonary hemorrhage. Acute PH is associated with robust recruitment of proinflammatory innate immune cells that disrupt the alveolar-capillary barrier^33,78^. In this context, the loss of ITK signaling significantly reduced the accumulation of CD11b⁺Ly6C⁺ inflammatory monocytes and CD11b⁺Ly6G⁺ neutrophils in the lung, and was also associated with reduced inflammatory pathology in the spleen and gastrointestinal tract. These findings suggest that ITK contributes broadly to the inflammatory cascade that drives both pulmonary and systemic tissue injury in PH. This tissue protection was paralleled by a marked suppression of systemic inflammatory cytokines. ITK deficiency significantly reduced circulating levels of IFN-γ, IL-6, TNF-α, and IL-17—mediators known to promote vascular permeability, inflammatory amplification, and tissue injury during severe autoimmune inflammation^19,79^. By limiting this proinflammatory milieu, modulation of the ITK axis may provide a dual therapeutic benefit: preserving alveolar-capillary barrier integrity while reducing the systemic inflammatory injury that contributes to poor outcomes in PH.

Beyond suppressing proinflammatory mediators, ITK deficiency also promoted a pro-resolving cytokine environment, including increased IL-10, IL-13, and, in some settings, IL-4^80,81^. This shift is particularly relevant in PH, where early containment of inflammatory injury is likely critical for preserving alveolar-capillary barrier integrity. These mediators are known to act not only on immune cells but also directly on the lung epithelium and endothelium. This observation suggests a dual-action mechanism in which Itk⁻/⁻Tregs suppress inflammatory infiltration while simultaneously signaling to the alveolar-capillary unit to enhance barrier integrity and promote tissue resilience. A central finding of our study is that the adoptive transfer of Itk⁻/⁻Tregs was sufficient to rescue established PH in WT recipients ^20,82^. Because the therapeutic efficacy of Tregs is often limited by impaired stability or function under inflammatory stress^83,84^, these findings suggest that the loss of ITK signaling supports a more durable regulatory program in vivo.

To define the mechanisms underlying this protective response, we examined the expression of Amphiregulin (AREG), a key regulator of tissue repair and immune stability^26,85–90^. We demonstrated that Tregs isolated from the lungs of Itk⁻/⁻mice express significantly higher levels of AREG, which actively protects mice from PH^91,92^. Functional relevance was further supported by our adoptive transfer experiments, in which Itk⁻/⁻ Tregs conferred robust protection, suppressing IFN-γ, TNF-α, and IL-17 while simultaneously enhancing IL-10 production (Fig. 6).

Our transcriptomic analysis provides a mechanistic framework for this phenotype. RNA sequencing revealed that ITK deficiency reprograms the Treg transcriptional landscape, with enrichment of pathways linked to oxidative phosphorylation (OXPHOS) ^93^, mTORC1^94,95^, STAT5 signaling^96,97^, and tissue repair. Together with the observed changes in glycolytic and cell cycle-associated programs, these data suggest that Itk⁻/⁻ Tregs acquire a state of enhanced metabolic fitness that may support their persistence and suppressive activity in inflamed tissue. Thus, the loss of ITK signaling appears to generate not simply more Tregs, but Tregs with molecular features consistent with heightened regulatory capacity.

Several limitations of the present study should be acknowledged. First, our transcriptional analyses were performed on Tregs isolated from unchallenged mice and therefore may not fully capture the molecular programs adopted by Tregs within the inflamed lung microenvironment following pristane-induced pulmonary injury. Future studies directly comparing lung-resident Tregs from pristane-treated wild-type and ITK-deficient mice will be important for defining the disease-specific mechanisms associated with protection. Second, because the current study utilized germline ITK-deficient mice, we cannot completely exclude contributions from ITK deficiency in other immune cell populations. Although our adoptive-transfer experiments demonstrate that ITK-deficient Tregs are sufficient to markedly reduce established pulmonary hemorrhage in wild-type recipients, lineage-specific conditional ITK-deficient models will be important for defining the Treg-intrinsic role of ITK signaling. These studies will further clarify the mechanistic and translational potential of targeting the ITK–Treg axis in inflammatory lung disease.

Collectively, these findings establish ITK as a promising therapeutic target in PH. Modulating the ITK axis may offer a strategy to suppress systemic inflammation while promoting tissue-protective immune regulation and the restoration of alveolar-capillary barrier integrity. More broadly, our study provides a mechanistic rationale for developing ITK-directed approaches in severe inflammatory lung disease. While our findings in the pristane-induced model are striking, further studies are required to determine whether pharmacological ITK inhibition can fully phenocopy the genetic deficiency observed here, and to evaluate the long-term stability of these reprogrammed Tregs in chronic models of systemic autoimmunity.

## Materials and Methods

### Mice

Both male and female mice were included as a biological variable in all studies, as previously described^35,98^. C57BL/6 mice were obtained from Charles River Laboratories or The Jackson Laboratory. All experiments were performed using mice 8–12 weeks of age, with age- and sex-matched controls included throughout. ITK⁻/⁻ mice and WT mice were bred onto the C57BL/6 background and crossed with the Foxp3tm1Flv/J strain, an X-linked knock-in reporter in which Foxp3-expressing cells are co-marked with monomeric red fluorescent protein (mRFP). In this model, mRFP faithfully reports endogenous Foxp3 expression. ITK⁻/⁻ and WT Foxp3-RFP mice were generously provided by Dr. Avery August (Cornell University)^14,17,19^.

### Reagents, cell lines, flow cytometry

Spleen or lung tissues were processed into single-cell suspensions. For surface staining, cells were incubated with an Fc receptor-blocking antibody (anti-CD16/32) followed by fluorochrome-conjugated monoclonal antibodies at a 1:100 dilution (unless otherwise specified) for 30 minutes at 4°C. Antibodies were purchased from BioLegend, eBioscience, or BD Pharmingen. Surface-staining antibodies included anti-mouse CD3 (BV605, 100237), CD4 (PE, 100408; BV510, 100559), CD8 (PE/Cy7, 100722; APC/Cy7, 140421), CD11b (FITC, 101205; PerCP-Cy5.5, 450112-82), CD25 (FITC, 101907; FITC, 101908; Pacific Blue, 102021), CD44 (Pacific Blue, 156006), CD62L (APC, 161218; APC/Cy7, 304814), CD122 (APC, 105912; APC, 123213), and Ly-6C (APC, 128015). PE anti-mouse Ly-6G Antibody Cat#27608 For transcription factor and intracellular cytokine staining, cells were fixed and permeabilized using the Foxp3/Transcription Factor Staining Buffer Set (eBioscience) according to the manufacturer’s protocol. Intracellular antibodies included Eomes (PE, 12-4875-80; PE/Cy7, 25-4877-42), Foxp3 (Pacific Blue, 126410), T-bet (FITC, 64481; BV421, 644816), TCF-1 (PE, 564217), IFN-γ (APC, 505810), and TNF-α (FITC, 502906). Dead cells were excluded using a LIVE/DEAD Fixable Viability Dye (BioLegend). Data were acquired on a BD LSRFortessa (BD Biosciences) and analyzed using FlowJo software (Tree Star), with singlets identified by forward- and side-scatter properties as previously described^19,58^.

### Consumables

All general laboratory consumables and plasticware were sourced from EIMMUNA Medical Supply.

### Proteinuria Assays

To assess renal involvement, urine was collected from mice at baseline (Day 0) and at specified intervals (Day 14 or Day 21) following pristane administration. Total urinary protein concentrations were quantified using the Pierce BCA Protein Assay Kit (Thermo Scientific, Cat# 23225) according to the manufacturer’s instructions. Briefly, urine samples were diluted 1:10 in PBS, and 25 µL of the diluted sample was combined with 200 µL of the BCA working reagent (50:1 ratio of Reagents A:B) in a 96-well microplate. The plate was briefly centrifuged (30 s at 1,250 rpm) to ensure uniform mixing and incubated at 37°C for 30 minutes. Absorbance was measured at 562 nm using a BioTek FLx800 plate reader. Protein concentrations (mg/mL) were determined using Gen5 software by interpolation from a bovine serum albumin (BSA) standard curve. Data were normalized to baseline values to account for physiological variation as previously shown by our lab and others^24^.

### Serum Cytokine Profiling

Systemic cytokine concentrations were assessed at baseline (day 0) and at the study endpoints (day 14 or day 21). Whole blood was collected via cardiac puncture, and serum was isolated by centrifugation. A comprehensive panel of cytokines—including IFN-γ, TNF-α, IL-5, IL-12, IL-6, IL-10, IL-9, IL-17A, IL-17F, IL-22, and IL-13—was quantified using the LEGENDplex™ Mouse Th Cytokine Panel (13-plex) (BioLegend, Cat# 741011) according to the manufacturer’s instructions^17,18^. Briefly, serum samples were incubated with capture beads, followed by detection antibodies and streptavidin-phycoerythrin (SA-PE). Data were acquired on a BD LSRFortessa (BD Biosciences). Absolute cytokine concentrations (pg/mL) were calculated using the LEGENDplex Data Analysis Software (BioLegend) by interpolation from a 5-parameter logistic (5PL) standard curve. All samples were analyzed in duplicate to ensure technical reproducibility, as previously described^17,18,24,32,58,99^.

### Histopathological Evaluation

Following euthanasia at day 14 or day 21 post-pristane challenge, systemic organs—including the lungs, liver, spleen, kidneys, and small intestine—were harvested and fixed in 10% neutral-buffered formalin for 24–48 hours. Tissues were processed for paraffin embedding, sectioned at 5 µm, and stained with hematoxylin and eosin (H&E) by the Histology Core Facility at SUNY Upstate Medical University. Histopathological evaluation, specifically focusing on the severity of PH, alveolar-capillary disruption, and systemic inflammatory infiltration, was performed by a board-certified pathologist (L.C.) who was blinded to the experimental groups. Tissue injury was graded according to previously established semi-quantitative criteria^24,100,101^. Representative images were captured using a light microscope, and histopathology scores were analyzed using the Mann-Whitney *U* test for non-parametric comparisons.

### Isolation and Purification of Regulatory T Cells

To facilitate the live isolation of regulatory T cells, we utilized Itk⁻/⁻ and WT Foxp3-RFP reporter mice on the C57BL/6 background. In this X-linked knock-in model, monomeric red fluorescent protein (mRFP) faithfully reports endogenous Foxp3 expression. Foxp3-RFP expression was confirmed in both WT and Itk⁻/⁻ cohorts prior to experimentation.

Single-cell suspensions were prepared from spleens, or lungs and CD4⁺ lymphocytes were pre-enriched using anti-CD4 magnetic microbeads and MACS column-based separation (Miltenyi Biotec, Cat# 130-114-043). For high-purity isolation, pre-enriched cells were stained with antibodies against CD3 and CD25. Canonical Tregs were identified and purified as CD3⁺CD4⁺CD25⁺Foxp3(RFP)⁺ cells, while non-canonical Tregs were identified as CD3⁺CD4⁺CD25⁻Foxp3(RFP)⁺ cells. Fluorescence-activated cell sorting (FACS) was performed on a BD FACSAria IIIu (BD Biosciences), with sorted Treg purity routinely exceeding 98%. Purified cells were used immediately for RNA sequencing or adoptive transfer. Unless otherwise noted, all cell culture reagents were obtained from Sigma-Aldrich or Invitrogen, consistent with previously established protocols^20,24,58^.

### Western blotting

For the immunoblotting data shown in Figure 5K, Tregs from either WT or Itk⁻/⁻ mice were isolated from lung tissue. Briefly, lungs were gently homogenized, and total lung cellularity was quantified. Lung CD4⁺ T cells were then enriched by MACS using CD4 microbeads (Miltenyi Biotec; Cat# 130-114-043). These cells were further purified by FACS for CD4⁺, CD25⁺, and Foxp3⁺ (via the RFP reporter). The sorted Tregs were lysed in freshly prepared RIPA buffer (Fisher Scientific; Cat# PI9900) supplemented with a complete protease inhibitor cocktail (Sigma-Aldrich; Cat# 11697498001) and clarified by centrifugation at 14,000 rpm for 10 minutes at 4°C. Lysates equivalent to 1 × 10⁶ cells per sample were resolved on 12%–18% SDS–polyacrylamide gels and transferred to nitrocellulose membranes for immunoblotting. Unless otherwise specified, reagents were obtained from Invitrogen (Grand Island, NY) or Sigma-Aldrich (St. Louis, MO). For analyses, membranes were probed with antibodies against Amphiregulin (AREG; clone AREG559, Thermo Fisher Scientific; Cat# 14-9999-82) and β-actin (Cell Signaling Technology; Cat# 4970), as previously described^24^.

### Statistics

Statistical analyses were performed using GraphPad Prism 10. Unless otherwise specified in the figure legends, data are presented as mean ± SD. For comparisons between two groups, an unpaired two-tailed Student’s *t*-test was used. For comparisons involving multiple groups, a one-way or two-way ANOVA was performed, followed by Tukey’s *post hoc* test for multiple comparisons. Non-parametric data, such as histopathology scores, were analyzed using the Mann-Whitney *U* test. A *p* value of ≤ 0.05 was considered statistically significant (\**p* ≤ 0.05, *p* ≤ 0.01, \*\*\**p* ≤ 0.001, \*\*\*\**p* ≤ 0.0001). For *in vivo* studies, cohorts of *n* = 10–22 age- and sex-matched mice were used per group. All experiments were independently repeated at least twice to ensure reproducibility. Sample sizes were determined based on power calculations to achieve a minimum power of 0.8, and the experimental design was consistent with previously established protocols^17–20,24,32,35,36,58,99,102–113^.

### RNA Sequencing and Transcriptomic Analysis

For transcriptomic profiling, canonical and non-canonical Tregs were FACS-purified from the spleens of WT (n = 3) and Itk*⁻/⁻* (n = 3) mice, as described above. Total RNA was extracted, and library preparation and high-throughput sequencing were performed by the Molecular Analysis Core Facility at SUNY Upstate Medical University. Raw sequencing data were processed in R (v4.5.1) using RStudio (v2025.05.1) and Bioconductor packages. Transcript abundance was quantified via pseudoalignment with Kallisto (v0.46.2) ^114^. Transcript abundance estimates were imported with tximport^115^, summarized to the gene level, and converted to scaled TPM values. Genes with low expression were filtered prior to model fitting. Differentially expressed genes (DEGs) were identified using DESeq2^116^, with significance defined using a Benjamini-Hochberg^117^, adjusted false discovery rate (FDR) ≤ 0.05 and a |log₂ fold change| ≥ 0.5. Variance-stabilized expression data were visualized using principal component analysis (PCA) and hierarchical clustering heatmaps generated with pheatmap^118^. Functional annotation of identified up- and downregulated genes was performed by Gene Ontology (GO) enrichment analysis using gprofiler2 (gost) ^119^. Gene Set Enrichment Analysis (GSEA) was conducted using clusterProfiler against the MSigDB Hallmark gene set, with the DESeq2 Wald statistic as the ranked input list. Normalized enrichment scores (NES) and ridge plots were used to identify metabolic and signaling pathways unique to ITK-deficient Treg subsets. All RNA-seq data have been deposited in the NCBI Gene Expression Omnibus (GEO) under accession number GSE185327.

### Ethics Statement and Animal Welfare

All animal housing and experimental procedures were performed in strict accordance with the recommendations in the *Guide for the Care and Use of Laboratory Animals* of the National Institutes of Health. The study protocol was reviewed and approved by the Institutional Animal Care and Use Committee (IACUC) of SUNY Upstate Medical University (Protocol #443). All mice were maintained in a pathogen-free facility under a 12-hour light/dark cycle with *ad libitum* access to food and water. Every effort was made to minimize animal suffering during the pristane challenge and subsequent procedures.

## Author contributions

M.S.H., H.X., A.M., I.H.T and R.T.: Investigation, methodology, and data acquisition. L.C. and L.S.: Formal analysis and histological evaluation. M.K.: Conceptualization, supervision, project administration, funding acquisition, resources, data curation, and writing—original draft preparation, review, and editing.

## Supporting information

Sup materai

## Acknowledgements

The authors thank all members of the Karimi Laboratory for their helpful discussions and technical support throughout this study. We also thank the Molecular Analysis Core and the Histology Core Facility at SUNY Upstate Medical University for their expertise and assistance with high-throughput sequencing and pathological processing.

## Copyright

2026, MSH et al. This is an open access article published under the terms of the Creative Commons Attribution 4.0 International License.

## Supplemental Figures

**Supplemental Figure 1.**
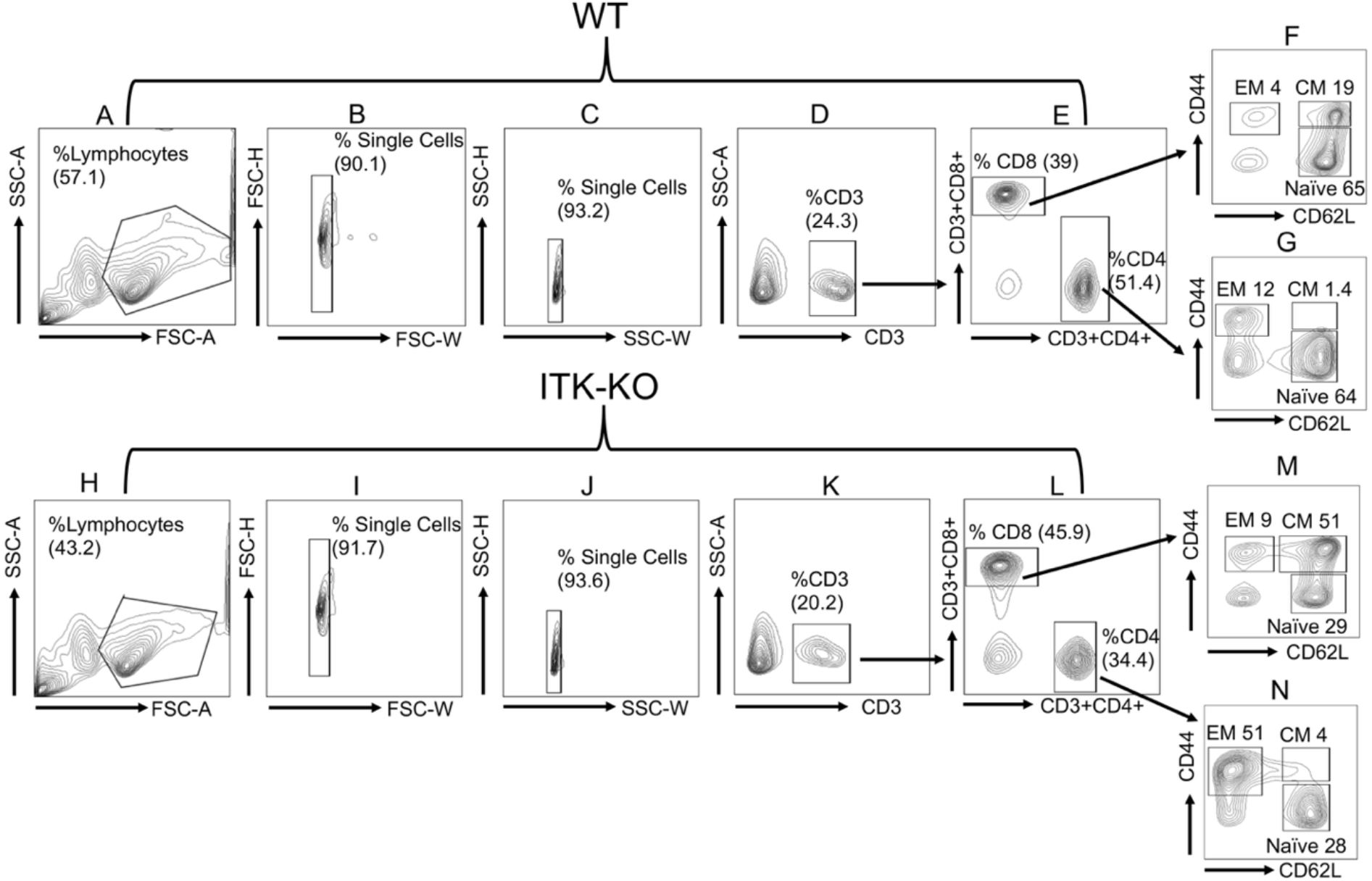
Gating strategies for effector and central memory phenotypes. Freshly isolated splenocytes from WT (A–G) and Itk⁻/⁻(H–N) mice were analyzed by flow cytometry. (A, H) Lymphocyte populations were initially gated based on side scatter area (SSC-A) versus forward scatter area (FSC-A). (B, I) Lymphocytes were then gated for single cells using forward scatter height (FSC-H) versus forward scatter width (FSC-W) to discriminate doublets. (C, J) Additional singlet gating was performed using side scatter height (SSC-H) versus side scatter width (SSC-W) to exclude any remaining doublets. (D, K) CD3⁺ T lymphocytes were identified from the single-cell population based on SSC-A versus CD3 expression. (E, L) The CD3⁺ compartment was further subdivided into CD8⁺ and CD4⁺ T cell populations. (F, M) CD8⁺ and (G, N) CD4⁺ T cells were then analyzed to identify and quantify effector memory (EM; CD44⁺CD62L⁻), central memory (CM; CD44⁺CD62L⁺), and naïve (CD44⁻CD62L⁺) subsets, as depicted in the contour plots.

**Supplemental Figure 2.**
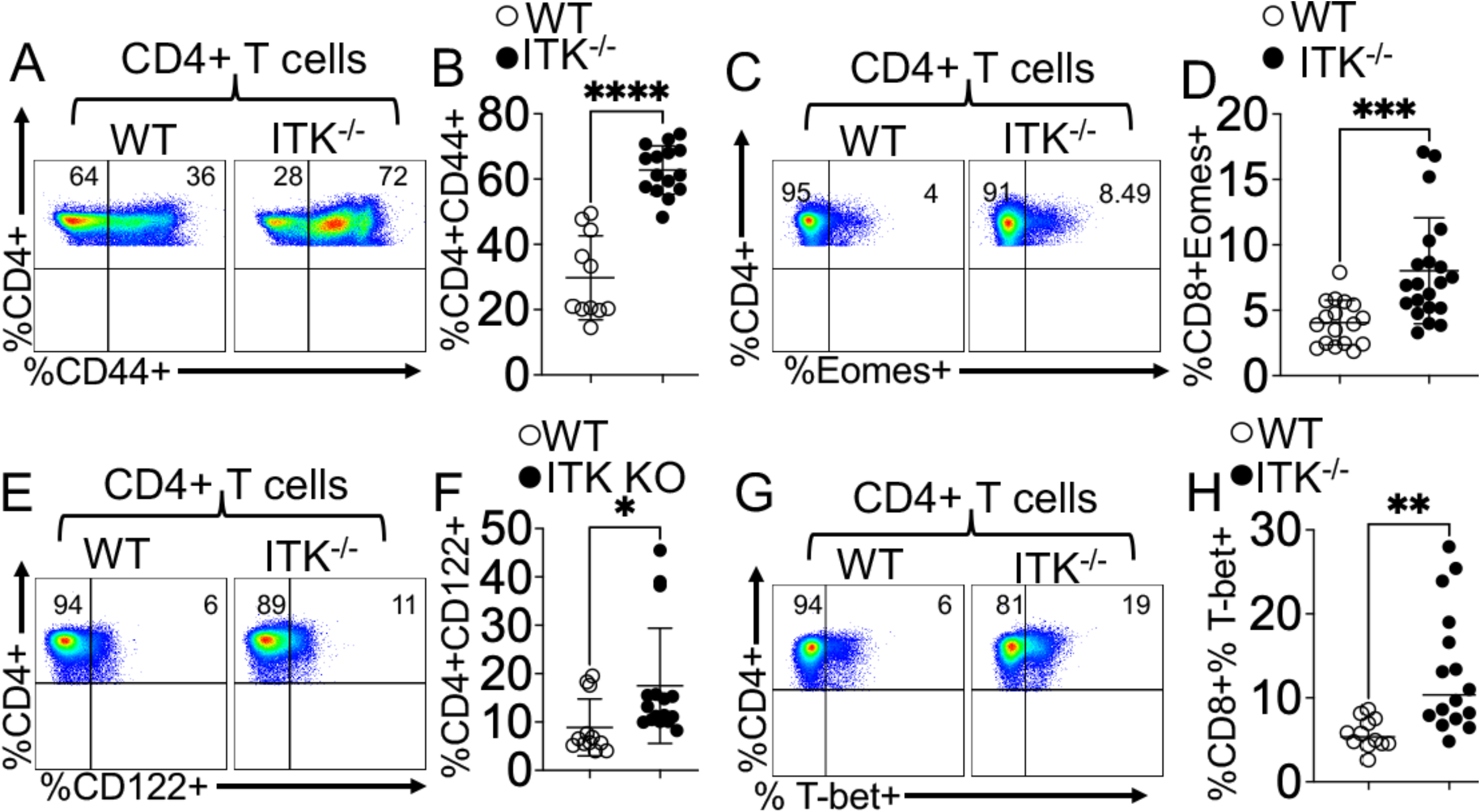
ITK deficiency promotes an activated, memory-like program in CD4⁺ T cells. (A and B) Surface expression of CD44 on splenic CD4⁺ T cells. Representative flow cytometry plots (A) and quantification of CD44⁺ frequencies (B) in WT and Itk⁻/⁻ mice. (C and D) Expression of the T-box transcription factor Eomes. Representative flow cytometry plots (C) and quantification of total Eomes⁺ CD4⁺ T cells (D) within the CD3⁺CD4⁺ gate. (E and F) Surface expression of CD122 (IL-2Rβ). Representative flow cytometry plots (E) and quantification of CD122 frequencies on CD4⁺ T cells (F) from WT and Itk⁻/⁻mice. (G and H) Intracellular expression of the transcription factor T-bet. Representative flow cytometry plots (G) and quantification of T-bet frequencies within the CD3⁺CD4⁺ population (H). Data are presented as mean ± SEM (*n* = 15–20 mice per group). Results are representative of at least 3 independent experiments. Statistical significance was determined by an unpaired two-tailed Student’s *t*-test \*\*\**p* < 0.001, * P ≤ 0.05 \*\*\*\**p* < 0.0001, ** P ≤ 0.01.

**Supplemental Figure 3.**
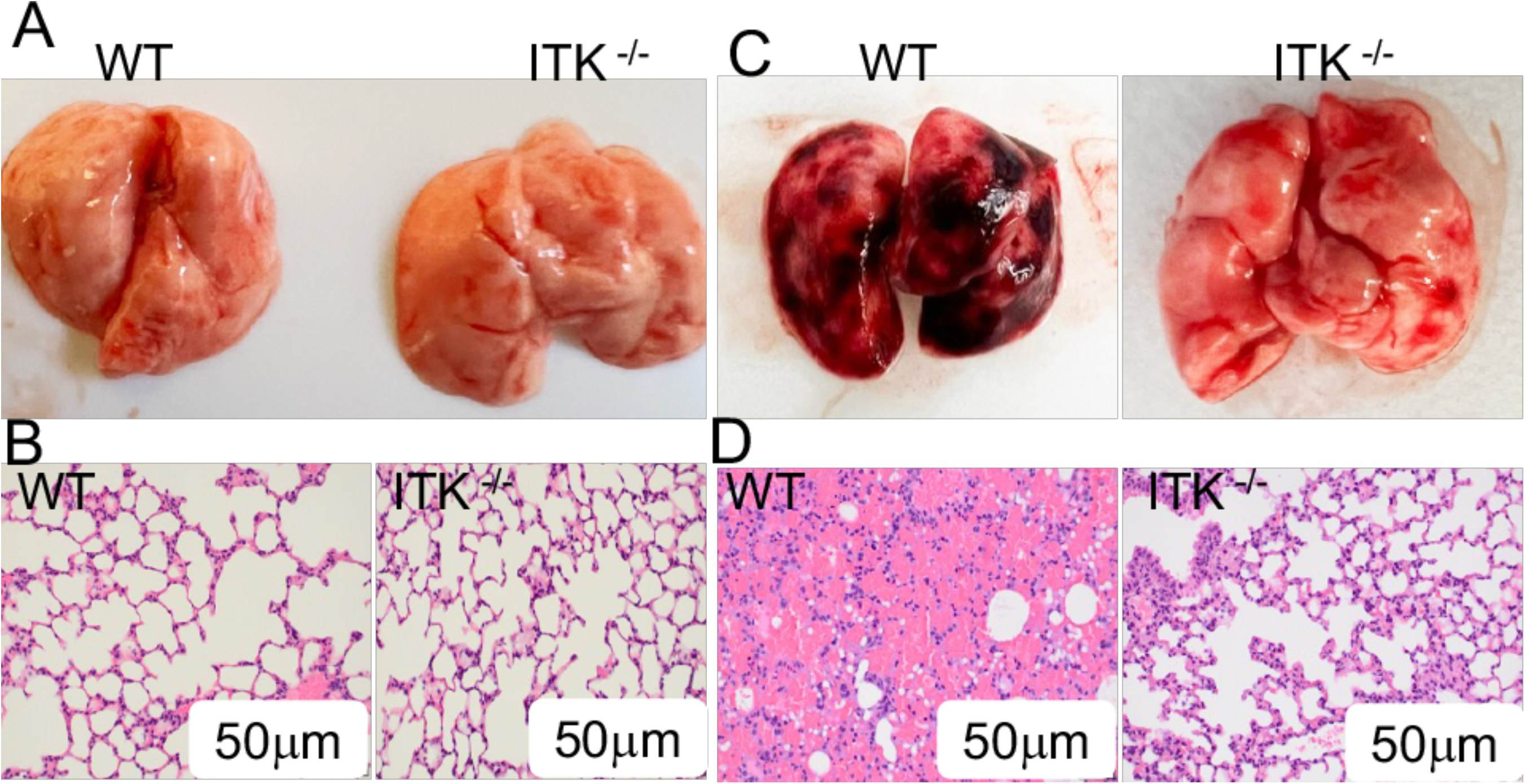
ITK deficiency protects mice from pristane-induced PH and systemic pathology. (A and B) Representative gross morphology of lungs and H&E-stained lung sections from WT and Itk⁻/⁻ mice without pristane injection. (C and D) Representative gross morphology and H&E-stained lung sections from WT and Itk⁻/⁻ mice 14 days after pristane injection

**Supplemental Figure 4.**
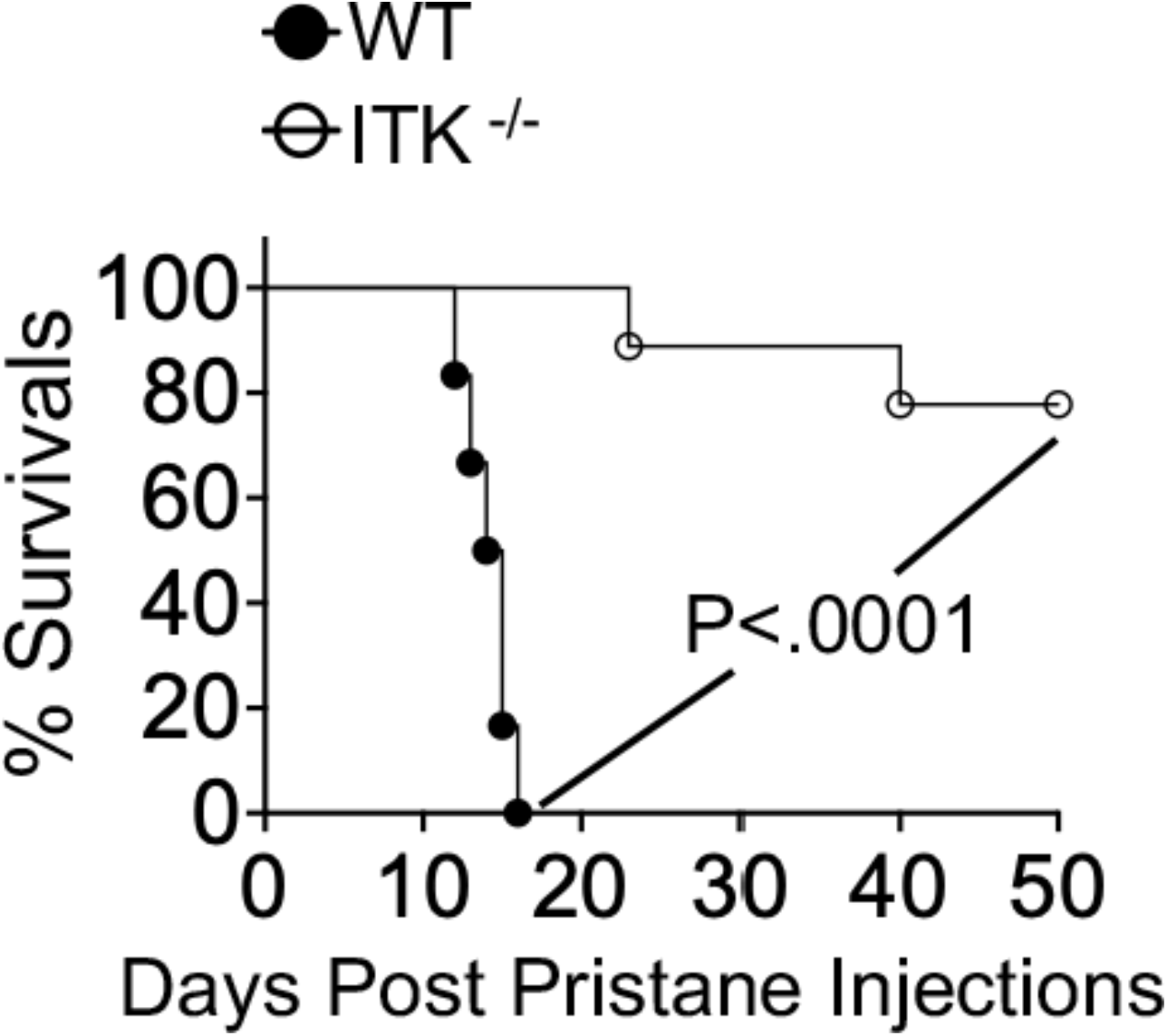
ITK deficiency prolongs survival following pristane-induced PH. WT and Itk⁻/⁻ mice received a single i.p. injection of 0.5 mL pristane to induce pulmonary hemorrhage (PH). Survival was monitored for 50 days post-injection.

**Supplemental Figure 5.**
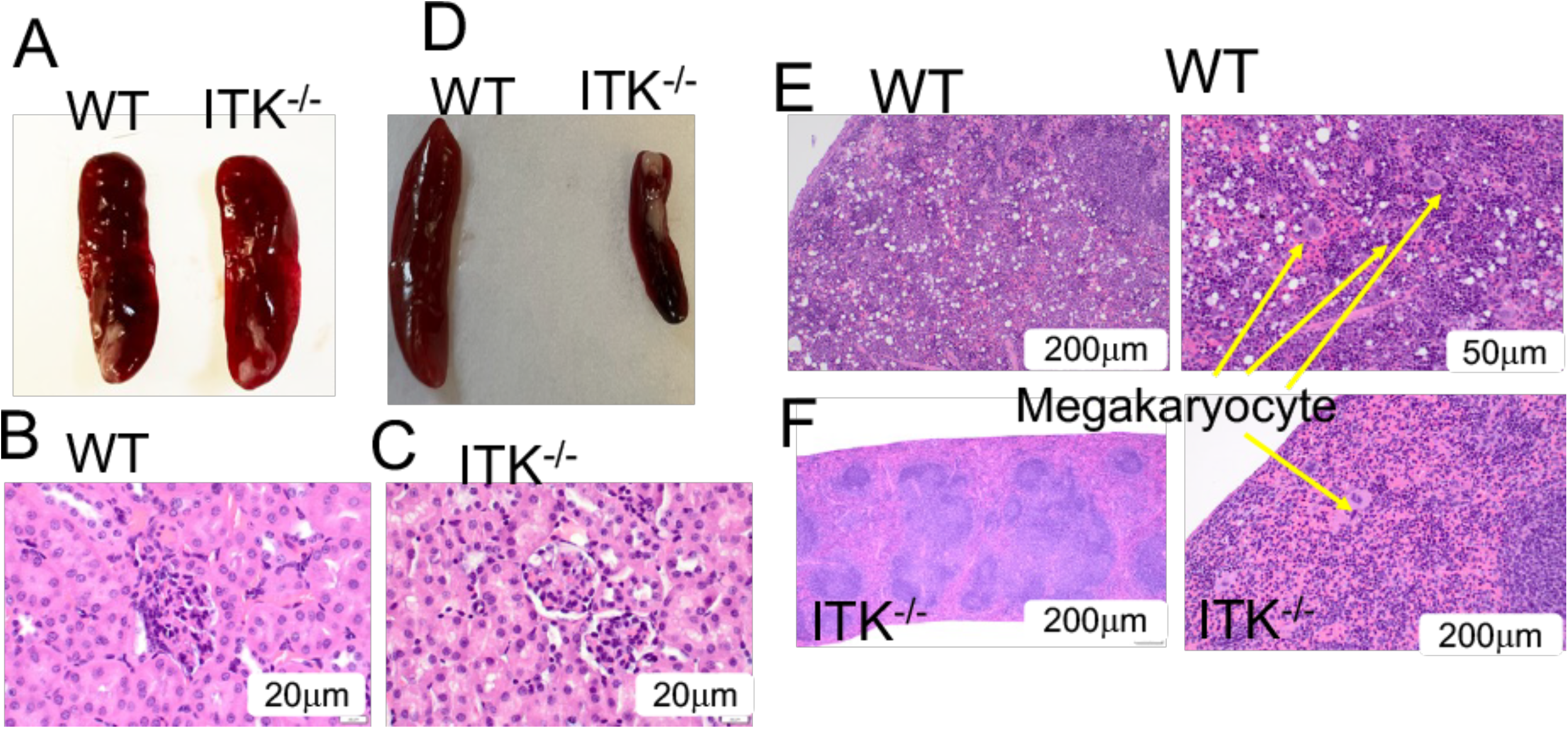
ITK deficiency protects mice from pristane-induced PH and systemic pathology. (A–C) Representative gross morphology of lungs and H&E-stained lung sections from either WT or Itk⁻/⁻ mice without pristane injection. (D–F) Representative gross images and H&E-stained sections of spleens from WT and Itk⁻/⁻ mice after pristane injection

**Supplemental Figure 6.**
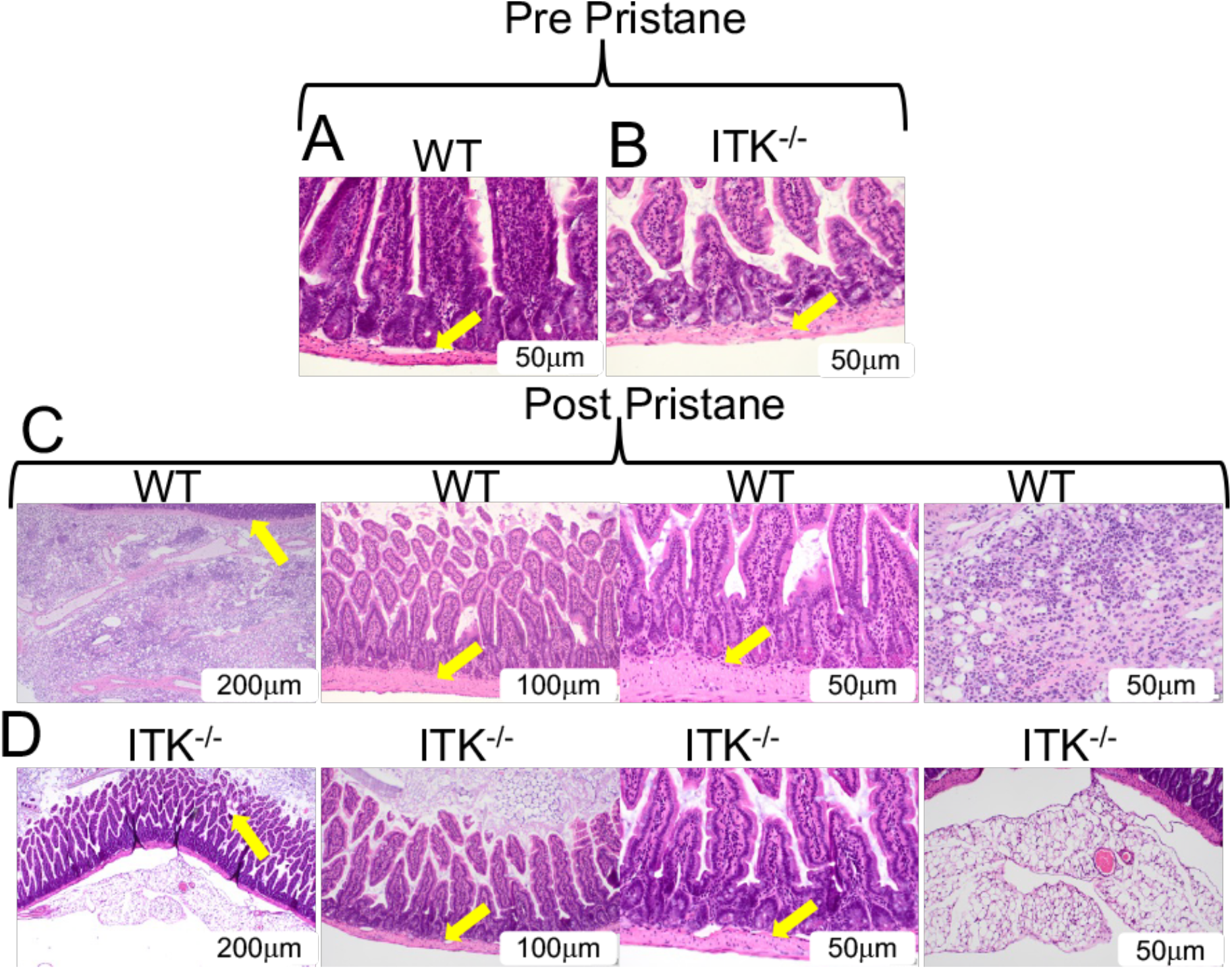
ITK deficiency protects mice from pristane-induced small intestine pathology. (A and B) Representative gross morphology and H&E-stained sections of the small intestine from either WT or Itk⁻/⁻ mice without pristane injection. (C and D) Histopathological evaluation of the small intestine. Representative H&E-stained sections of the small intestine from WT (C) and Itk⁻/⁻ (D) mice at day 14 post-pristane administration. In WT mice, systemic inflammatory injury is associated with evident mucosal damage, characterized by villus architecture disruption and increased inflammatory cell infiltration (black arrows). In contrast, Itk⁻/⁻mice exhibited significant protection against this systemic injury, with preservation of mucosal integrity. Results are representative of 3 independent experiments.

**Supplemental Figure 7.**
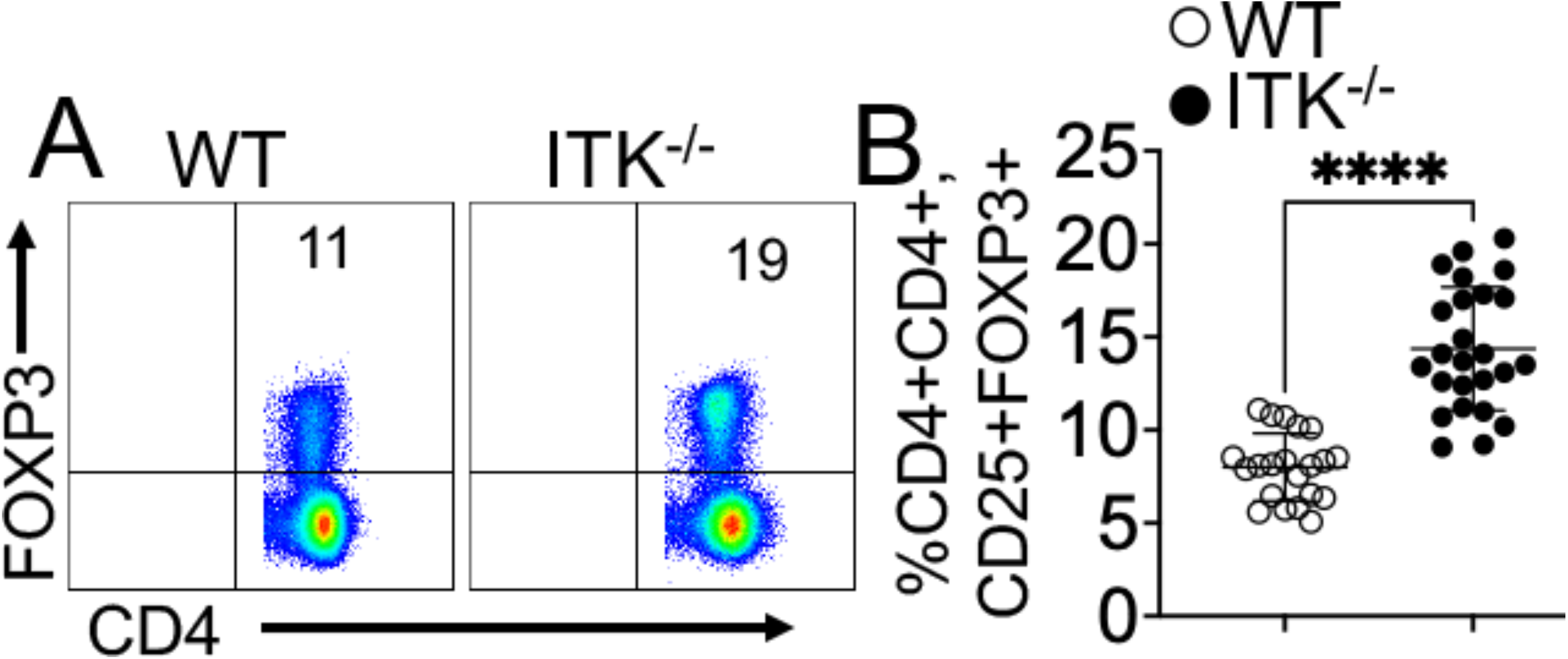
ITK deficiency promotes the expansion of non-canonical Tregs. (A and B) Quantitative analysis of Treg subsets at baseline. Representative flow cytometry plots (A) and statistical quantification (B) of freshly isolated splenic regulatory T cells from WT and Itk⁻/⁻ mice. Tregs were identified as CD3⁺CD4⁺Foxp3⁺ cells. ITK deficiency led to a significant increase in the frequency and absolute number of non-canonical (CD25⁻) Tregs compared with WT controls, while maintaining the canonical (CD25⁺) population. This expansion contributes to the broader regulatory reservoir observed in Itk⁻/⁻mice. Data are presented as mean ± SEM (*n* = 15–20 mice per group). Results are representative of 3 independent experiments. Statistical significance was determined by an unpaired two-tailed Student’s *t*-test. \*\*\*\**p* ≤ 0.0001.

**Supplemental Figure 8.**
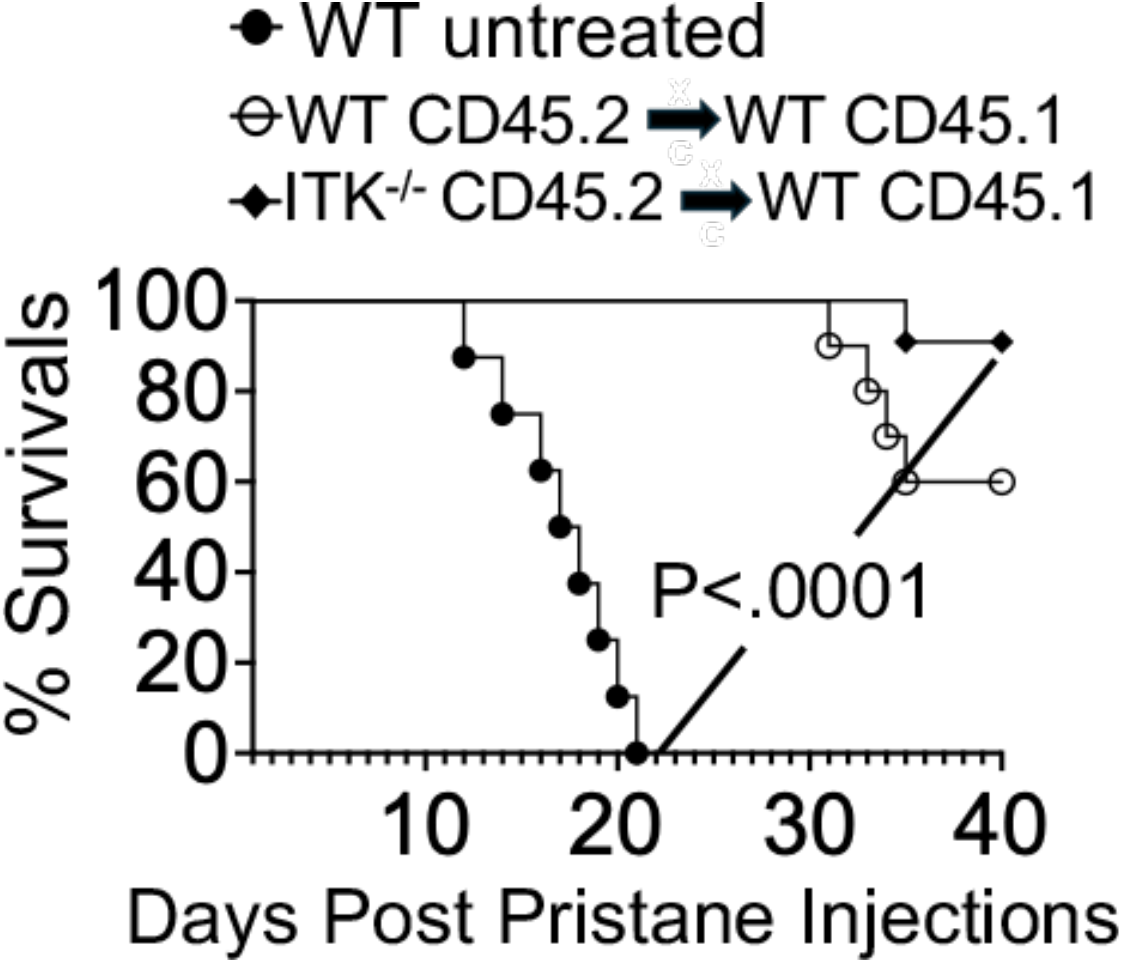
Adoptive transfer of Itk⁻/⁻ Tregs prolongs survival following pristane-induced PH. WT host mice (CD45.1⁺) received a single i.p. injection of 0.5 mL pristane. Mice were divided into three groups: left untreated, or adoptively transferred with Tregs isolated from either WT or Itk⁻/⁻ donor mice (CD45.2⁺). Survival was monitored for over 40 days post-injection.

**Supplemental Figure 9.**
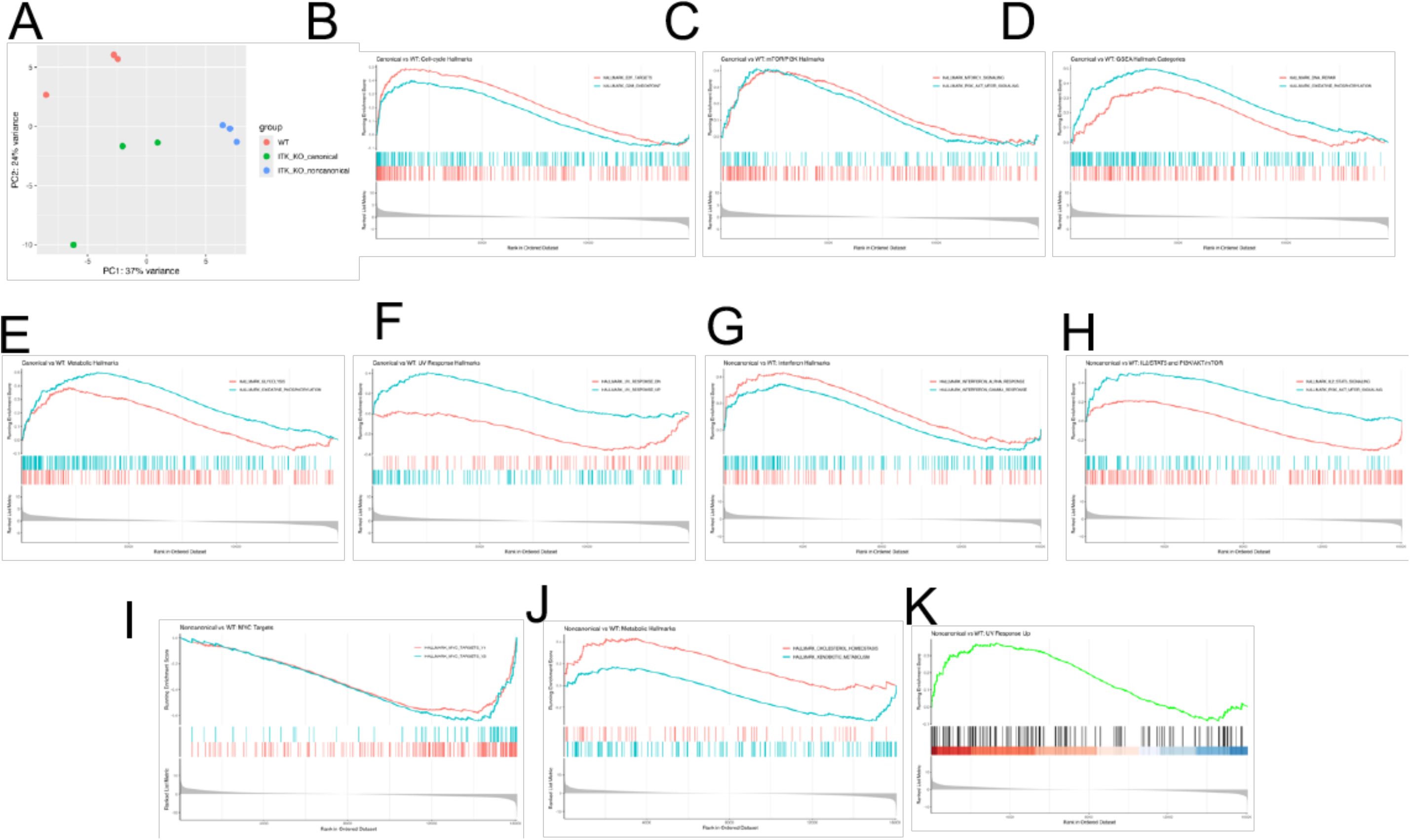
ITK Deficiency Reprograms Regulatory T Cells through Enhanced Metabolic and Signaling Circuitry. (A–J) Enrichment of metabolic and survival pathways. Summary of MSigDB Hallmark pathway enrichment in canonical and non-canonical ITK⁻/⁻ Tregs relative to WT controls. GSEA enrichment score plots show running enrichment scores for selected Hallmark gene sets. Representative GSEA plots are shown for high-impact pathways, including cell cycle progression, mTORC1 signaling, PI3K/Akt/mTOR signaling, IL-2/STAT5 signaling, MYC targets, and oxidative phosphorylation. Notably, ITK⁻/⁻ Tregs exhibit coordinated upregulation of metabolic, survival, and stress-adaptation pathways, including interferon response and UV response/DNA repair signatures, consistent with enhanced resilience in inflammatory environments. Data are derived from transcriptomic analysis of n = 3 independent biological replicates per group.

