## Supplementary material for "ITK deficiency protects against autoimmune pulmonary hemorrhage by promoting Amphiregulin-producing regulatory T cells": Sup materai

### Supplemental Figures

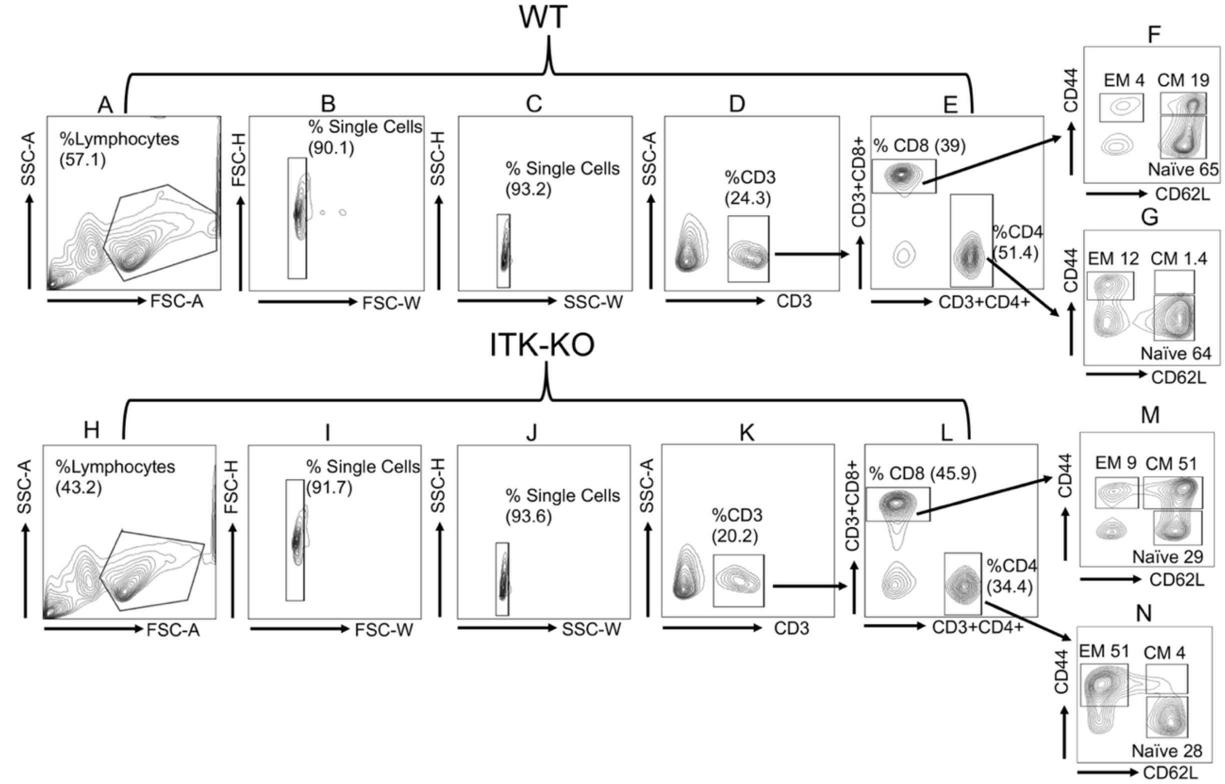


**Supplemental Figure 1. Gating strategies for effector and central memory phenotypes.** Freshly isolated splenocytes from WT (A–G) and Itk$^{-/-}$ (H–N) mice were analyzed by flow cytometry. (A, H) Lymphocyte populations were initially gated based on side scatter area (SSC-A) versus forward scatter area (FSC-A). (B, I) Lymphocytes were then gated for single cells using forward scatter height (FSC-H) versus forward scatter width (FSC-W) to discriminate doublets. (C, J) Additional singlet gating was performed using side scatter height (SSC-H) versus side scatter width (SSC-W) to exclude any remaining doublets. (D, K) CD3$^{+}$ T lymphocytes were identified from the single-cell population based on SSC-A versus CD3 expression. (E, L) The CD3$^{+}$ compartment was further subdivided into CD8$^{+}$ and CD4$^{+}$ T cell populations. (F, M) CD8$^{+}$ and (G, N) CD4$^{+}$ T cells were then analyzed to identify and quantify effector memory (EM; CD44$^{+}$CD62L$^{-}$), central memory (CM; CD44$^{+}$CD62L$^{+}$), and naïve (CD44$^{-}$CD62L$^{+}$) subsets, as depicted in the contour plots.

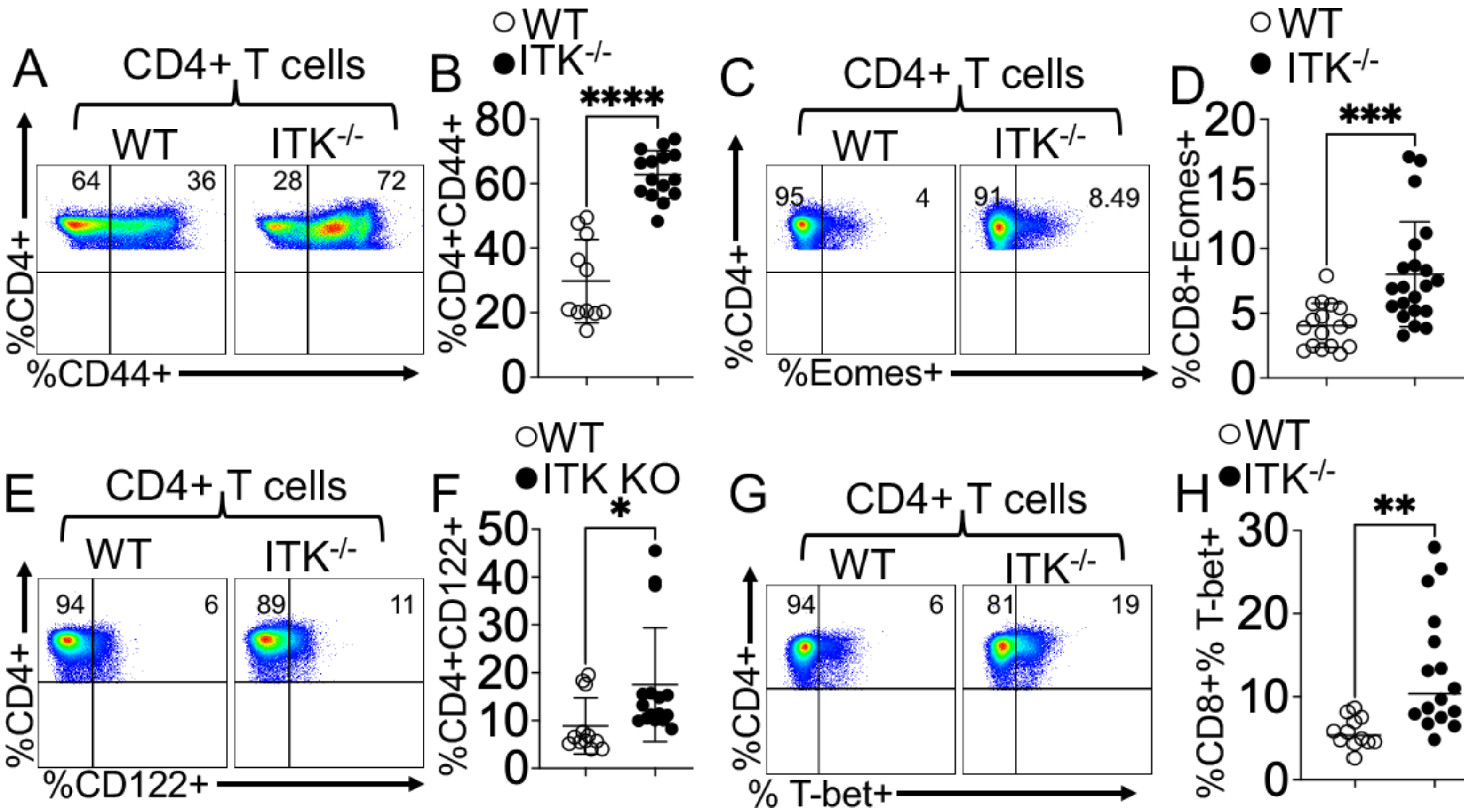


**Supplemental Figure 2. ITK deficiency promotes an activated, memory-like program in CD4$^+$ T cells.** (A and B) Surface expression of CD44 on splenic CD4$^+$ T cells. Representative flow cytometry plots (A) and quantification of CD44$^+$ frequencies (B) in WT and Itk$^{-/-}$ mice. (C and D) Expression of the T-box transcription factor Eomes. Representative flow cytometry plots (C) and quantification of total Eomes$^+$ CD4$^+$ T cells (D) within the CD3$^+$CD4$^+$ gate. (E and F) Surface expression of CD122 (IL-2Rβ). Representative flow cytometry plots (E) and quantification of CD122 frequencies on CD4$^+$ T cells (F) from WT and Itk$^{-/-}$ mice. (G and H) Intracellular expression of the transcription factor T-bet. Representative flow cytometry plots (G) and quantification of T-bet frequencies within the CD3$^+$CD4$^+$ population (H). Data are presented as mean ± SEM ($n$ = 15–20 mice per group). Results are representative of at least 3 independent experiments. Statistical significance was determined by an unpaired two-tailed Student's $t$-test ***$p < 0.001$, * $P \leq 0.05$ ****$p < 0.0001$, ** $P \leq 0.01$.

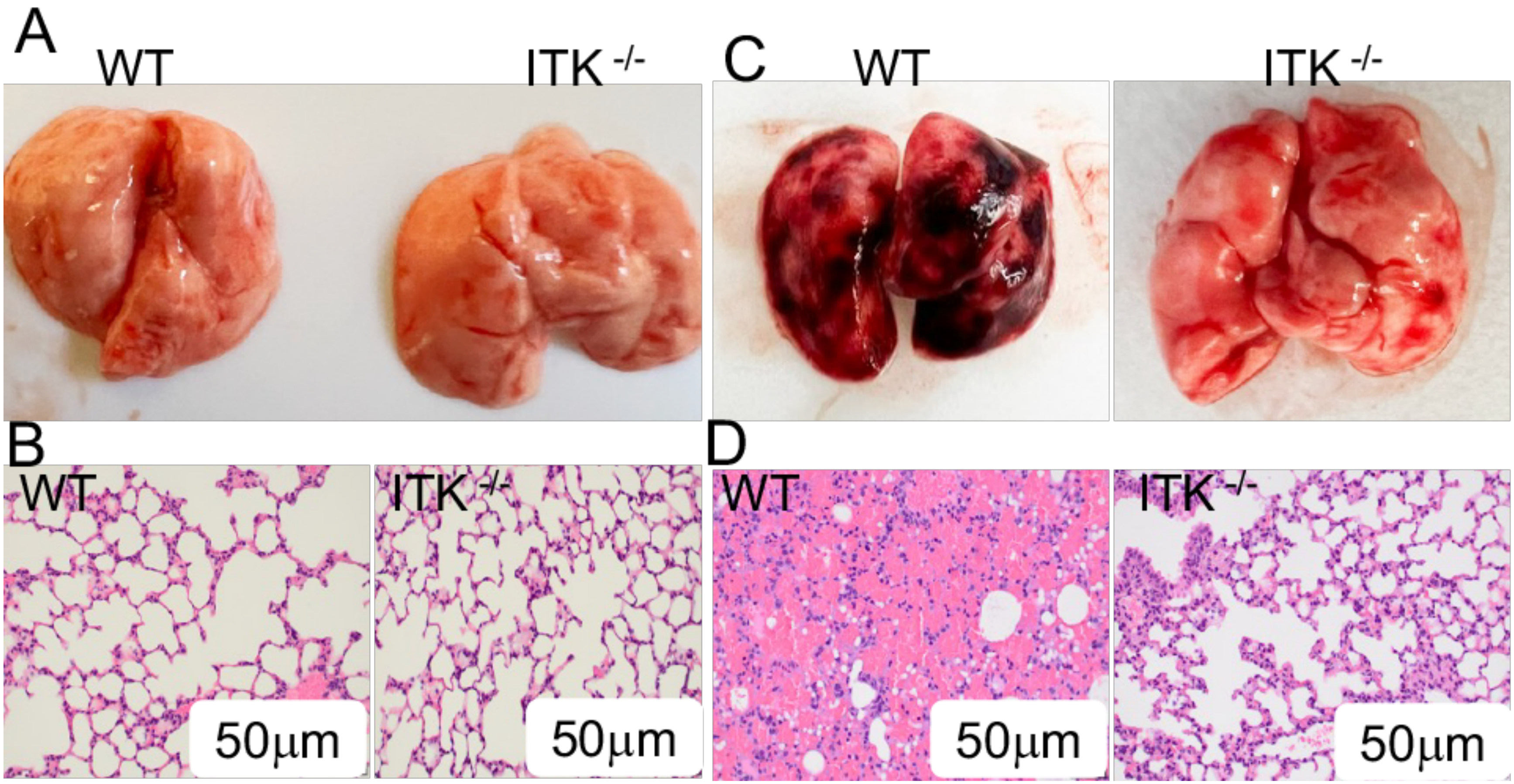


**Supplemental Figure 3. ITK deficiency protects mice from pristane-induced PH and systemic pathology.** (A and B) Representative gross morphology of lungs and H&E-stained lung sections from WT and Itk$^{-/-}$ mice without pristane injection. (C and D) Representative gross morphology and H&E-stained lung sections from WT and Itk$^{-/-}$ mice 14 days after pristane injection

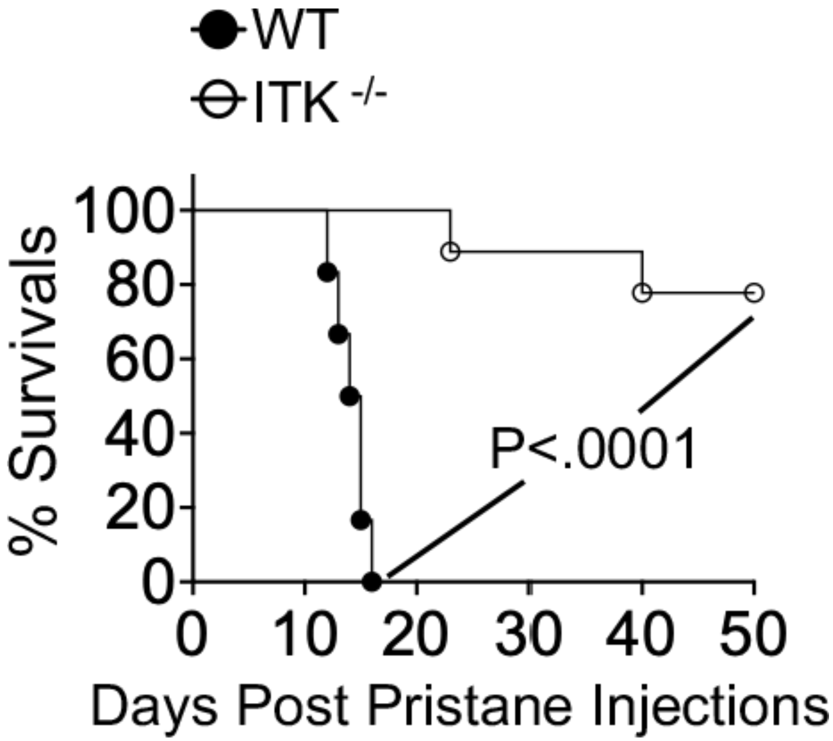


**Supplemental Figure 4. ITK deficiency prolongs survival following pristane-induced PH.** WT and Itk$^{-/-}$ mice received a single i.p. injection of 0.5 mL pristane to induce pulmonary hemorrhage (PH). Survival was monitored for 50 days post-injection.

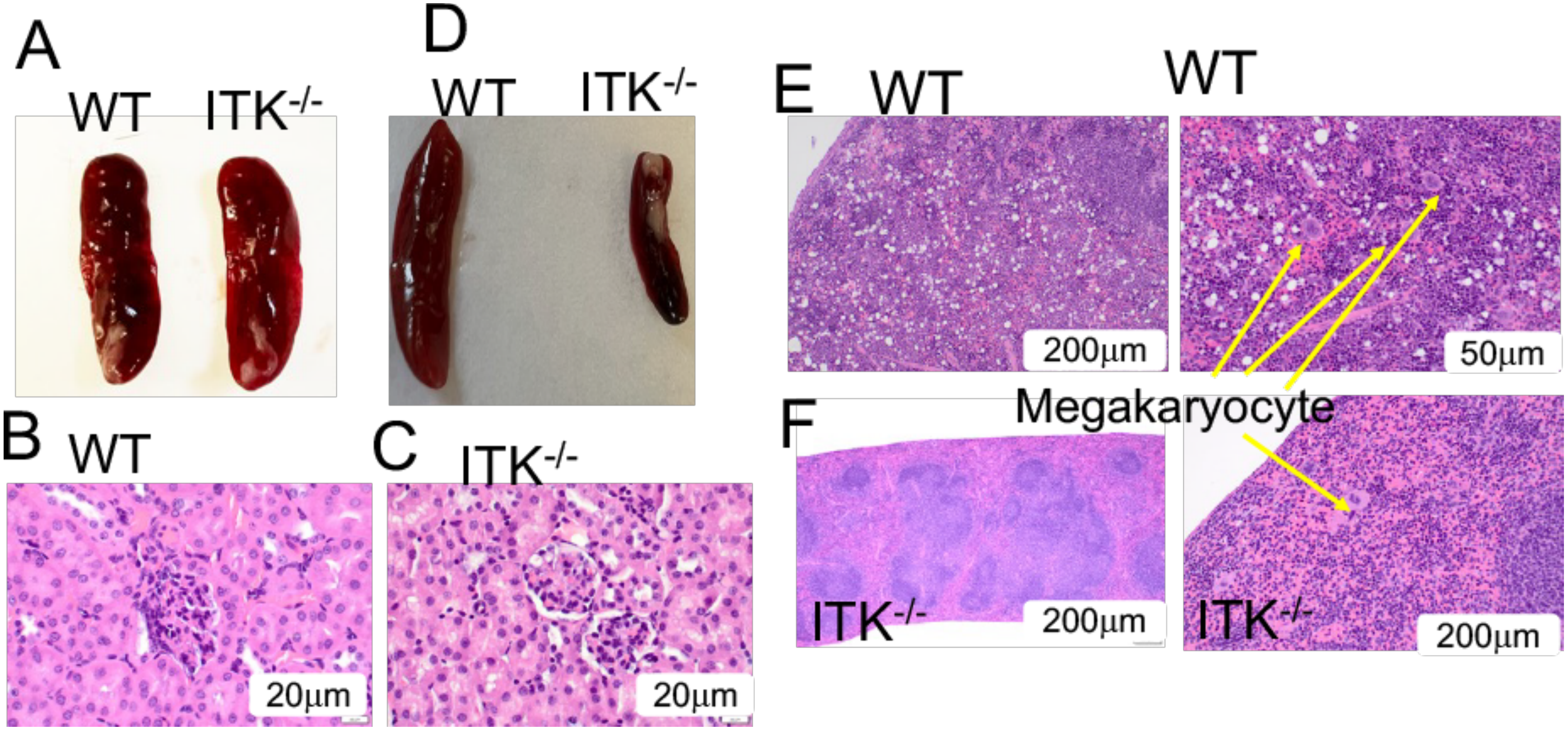


**Supplemental Figure 5. ITK deficiency protects mice from pristane-induced PH and systemic pathology.** (A–C) Representative gross morphology of lungs and H&E-stained lung sections from either WT or Itk$^{-/-}$ mice without pristane injection. (D–F) Representative gross images and H&E-stained sections of spleens from WT and Itk$^{-/-}$ mice after pristane injection

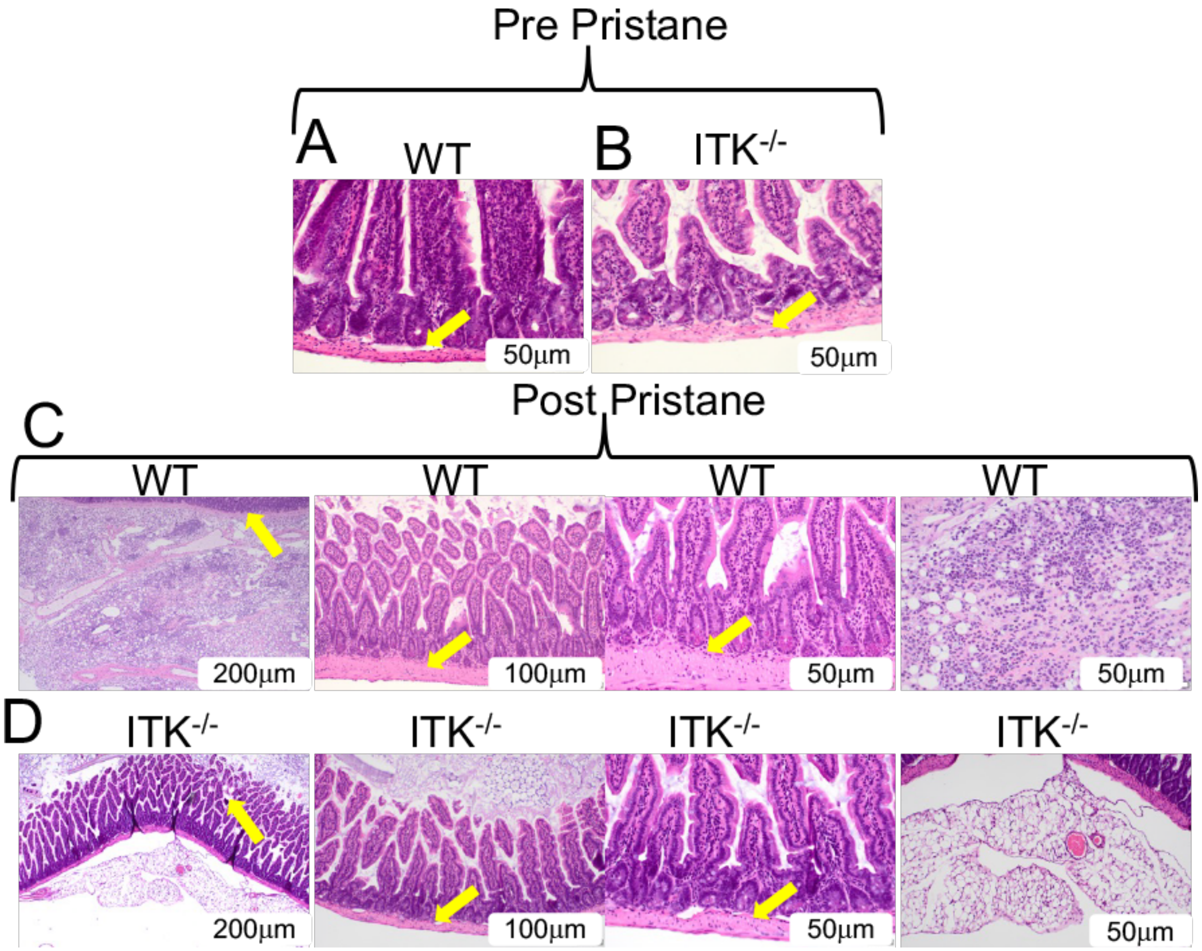


**Supplemental Figure 6. ITK deficiency protects mice from pristane-induced small intestine pathology.** (A and B) Representative gross morphology and H&E-stained sections of the small intestine from either WT or $Itk^{-/-}$ mice without pristane injection. (C and D) Histopathological evaluation of the small intestine. Representative H&E-stained sections of the small intestine from WT (C) and $Itk^{-/-}$ (D) mice at day 14 post-pristane administration. In WT mice, systemic inflammatory injury is associated with evident mucosal damage, characterized by villus architecture disruption and increased inflammatory cell infiltration (black arrows). In contrast, $Itk^{-/-}$ mice exhibited significant protection against this systemic injury, with preservation of mucosal integrity. Results are representative of 3 independent experiments.

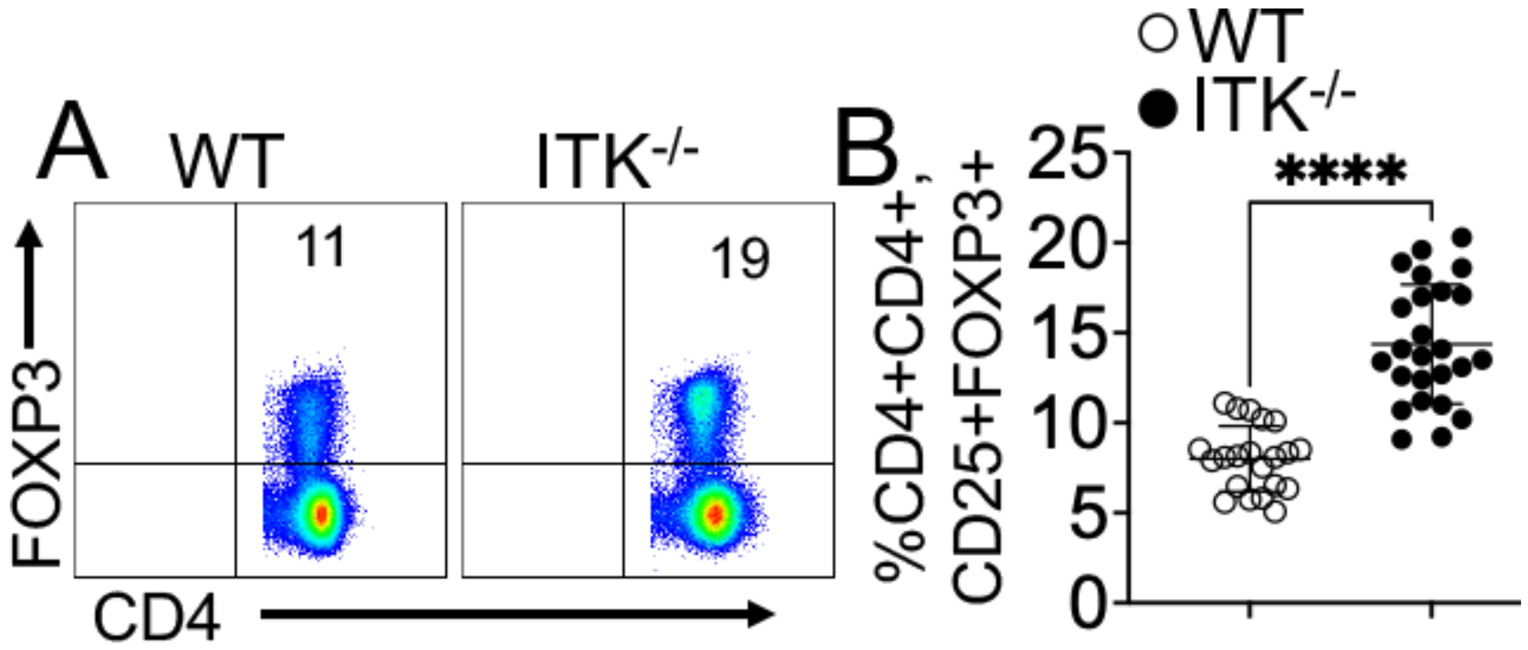


**Supplemental Figure 7. ITK deficiency promotes the expansion of non-canonical Tregs.** (A and B) Quantitative analysis of Treg subsets at baseline. Representative flow cytometry plots (A) and statistical quantification (B) of freshly isolated splenic regulatory T cells from WT and $Itk^{-/-}$ mice. Tregs were identified as $CD3^{+}CD4^{+}Foxp3^{+}$ cells. ITK deficiency led to a significant increase in the frequency and absolute number of non-canonical ($CD25^{-}$) Tregs compared with WT controls, while maintaining the canonical ($CD25^{+}$) population. This expansion contributes to the broader regulatory reservoir observed in $Itk^{-/-}$ mice. Data are presented as mean ± SEM ($n$ = 15–20 mice per group). Results are representative of 3 independent experiments. Statistical significance was determined by an unpaired two-tailed Student's $t$-test. ****$p \leq 0.0001$.

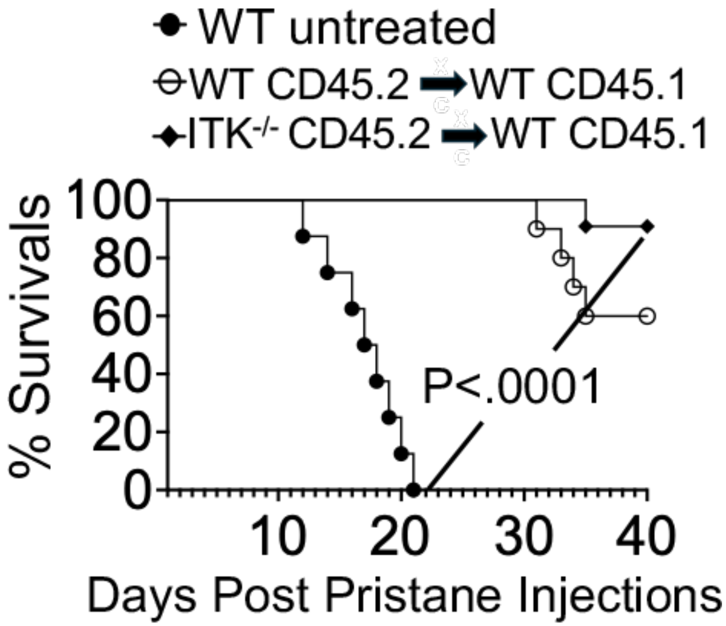


**Supplemental Figure 8. Adoptive transfer of Itk$^{-/-}$ Tregs prolongs survival following pristane-induced PH.** WT host mice (CD45.1$^{+}$) received a single i.p. injection of 0.5 mL pristane. Mice were divided into three groups: left untreated, or adoptively transferred with Tregs isolated from either WT or Itk$^{-/-}$ donor mice (CD45.2$^{+}$). Survival was monitored for over 40 days post-injection.

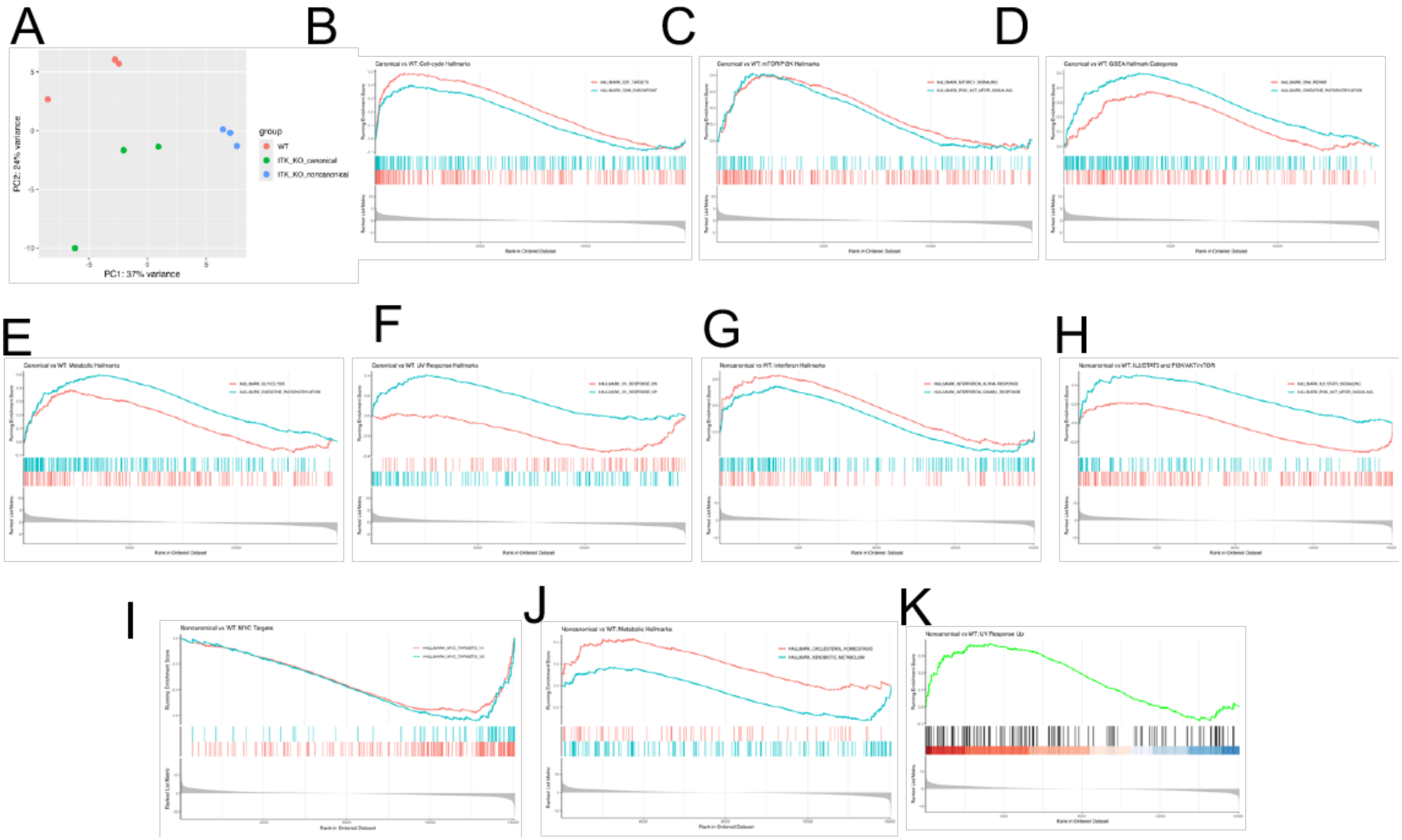
A
PC2: 24% variance
PC1: 37% variance
group
WT
ITK_KO_canonical
ITK_KO_noncanonical
B
Canonical vs WT: Cell-cycle Hallmarks
C
Canonical vs WT: mTOR/PI3K Hallmarks
D
Canonical vs WT: GSEA Hallmark Categories
E
Canonical vs WT: Metabolic Hallmarks
F
Canonical vs WT: UV Response Hallmarks
G
Noncanonical vs WT: Interferon Hallmarks
H
Noncanonical vs WT: IL2/STAT5 and PI3K/AKT/mTOR
I
Noncanonical vs WT: MYC Targets
J
Noncanonical vs WT: Metabolic Hallmarks
HALLMARK_CHOLESTEROL_HOMEOSTASIS
HALLMARK_XENOBIOTIC_METABOLISM
K
Noncanonical vs WT: UV Response Up

**Supplemental Figure 9. ITK Deficiency Reprograms Regulatory T Cells through Enhanced Metabolic and Signaling Circuitry.** (A–J) Enrichment of metabolic and survival pathways. Summary of MSigDB Hallmark pathway enrichment in canonical and non-canonical ITK$^{-/-}$ Tregs relative to WT controls. GSEA enrichment score plots show running enrichment scores for selected Hallmark gene sets. Representative GSEA plots are shown for high-impact pathways, including cell cycle progression, mTORC1 signaling, PI3K/Akt/mTOR signaling, IL-2/STAT5 signaling, MYC targets, and oxidative phosphorylation. Notably, ITK$^{-/-}$ Tregs exhibit coordinated upregulation of metabolic, survival, and stress-adaptation pathways, including interferon response and UV response/DNA repair signatures, consistent with enhanced resilience in inflammatory environments. Data are derived from transcriptomic analysis of n = 3 independent biological replicates per group.
